# Plexin/Semaphorin Antagonism Orchestrates Collective Cell Migration, Gap Closure and Organ sculpting by Contact-Mesenchymalization

**DOI:** 10.1101/2024.10.10.617649

**Authors:** Maik C. Bischoff, Jenevieve E. Norton, Mark Peifer

## Abstract

Cell behavior emerges from the intracellular distribution of properties like protrusion, contractility and adhesion. Thus, characteristic emergent rules of collective migration can arise from cell-cell contacts locally tweaking architecture – orchestrating self-regulation during development, wound healing, and cancer progression. The new *Drosophila* testis-nascent-myotube-system allows dissection of contact-dependent migration in vivo at high resolution. Here, we describe a process driving gap-closure during migration: Contact-mesenchymalization via the axon guidance factor Plexin A. This is crucial for testis myotubes to migrate as a continuous sheet, allowing normal sculpting-morphogenesis. Cells must stay filopodial and dynamically ECM-tethered near cell-cell contacts to spread while collectively moving. Our data suggest Semaphorin 1B acts as a Plexin A antagonist, fine-tuning activation. Our data reveal a contact-dependent mechanism to maintain sheet-integrity during migration, driving organ-morphogenesis using a highly conserved pathway. This is relevant for understanding mesenchymal organ-sculpting and gap-closure in migratory contexts like angiogenesis.

## Introduction

Collective cell migration is crucial for morphogenesis, regeneration, and cancer metastasis[1–3]. Migrating cells are often viewed as mesenchymal, but diverse models emphasize that the epithelial-to-mesenchymal transition (EMT) is not a binary switch. Instead, cells reside on a continuum and individual cells can even differ in this way locally. For example, migrating endothelia can have junctional lamellipodia and filopodia, while local mesenchymal dynamics can support epithelial intercalation, wound closure, and extrusion[4–8]. Often, mesenchymal cells self-regulate migration, with cell interactions enforcing or replacing external directional cues[9]. Cell-cell contact plays a very important role, creating characteristic emergent rules of locomotion by tweaking local architecture. The best-described example is contact inhibition of locomotion (CIL), but there are other “rule-sets” like contact-stimulation of migration or contact-following[10]. Understanding how changes in local architecture upon contact enable diverse emergent collective dynamics is a key challenge for the field.

Recently we established a new model system for contact-regulated collective cell migration in *Drosophila,* allowing us to combine the powerful genetic toolkit with 4D-live cell imaging at a level close to that in 2D-cell culture[11, 12]. Testis nascent myotube (TNM) collective cell migration encloses the pupal testis in the cells that assemble this organ’s characteristic circumferential musculature. After myoblast fusion, TNM migrate from the genital disc onto the testis, moving into a tightly confined space between overlying pigment cells and underlying cyst cells (Fig 1a)[13–15]. State-of-the-art microscopy allows us to live-image migration, track single cells, and mathematically analyze migration trajectories (Fig 1a’, Video S1)[11]. Each TNM in the sheet is highly protrusive, even at cell-cell contacts with interdigitating filopodial N-Cadherin junctions. Our previous work suggested directionality is a consequence of contact-dependent downregulation of matrix-adhesion stability, while free-edge filopodia undergo a normal molecular clutch-cycle[11]. Thus, directionality does not require an external polarizing cue. Notably, short-lived integrin adhesions are still present near mesenchymal cell-cell contacts, a finding we will address here.

**Figure 1.**
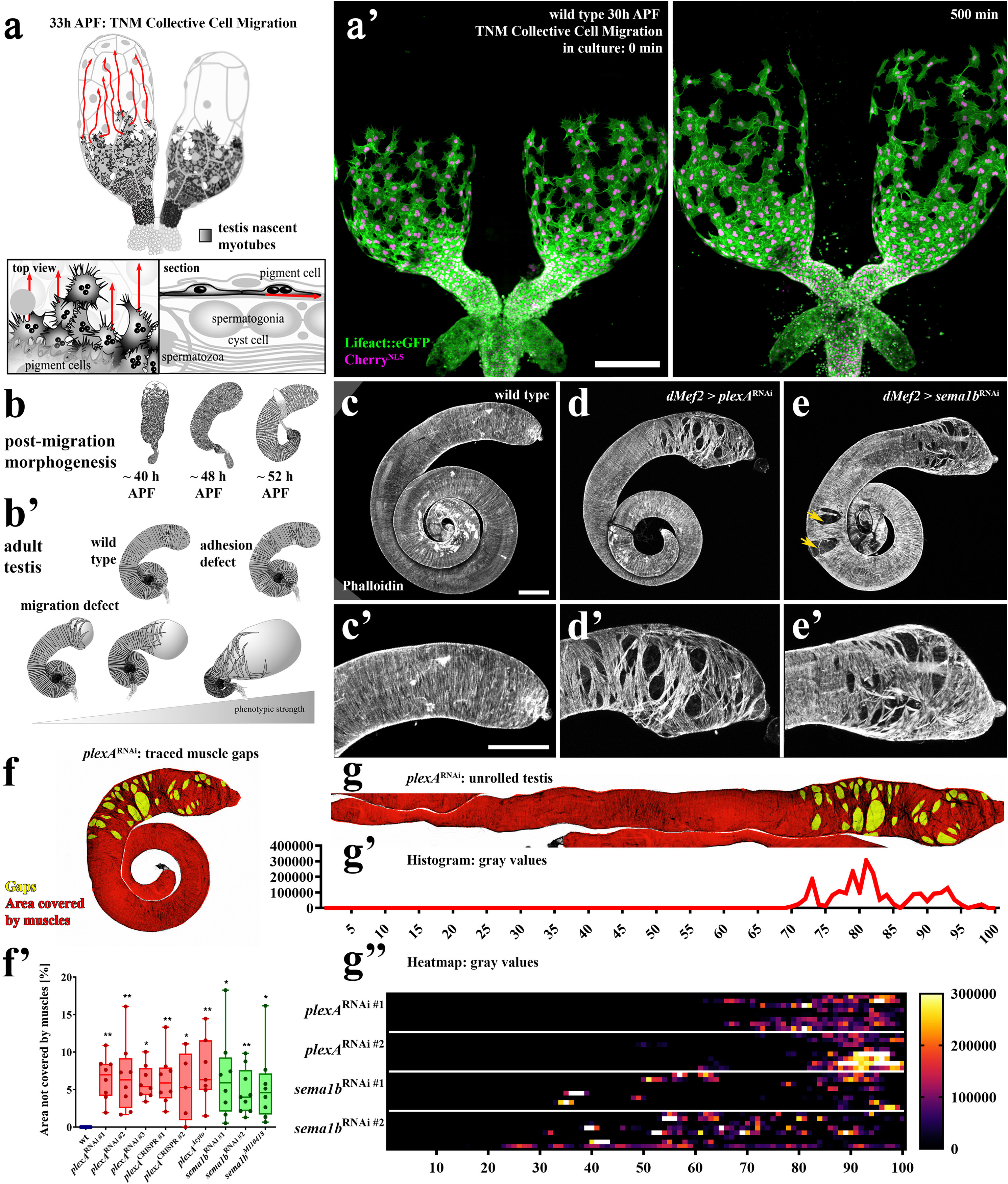
*plexA* and *sema1B* knockdown cause gap formation defects in the testis musculature suggesting migration defects. (a) Schematics of TNM collective cell migration during pupal development (a’) Live imaging of ex-vivo culture of testis with migrating TNM expressing *Lifeact::eGFP* and *Cherry^nls^* under the control of *dMef2-Gal4*. Scale bars =100 µm. (b) Steps of muscle-dependent testis morphogenesis after migration at 25°C. (b’) Schematic of types of adult muscle-dependent morphogenetic testis defects. (c-e) Adult testis with musculature stained with Phalloidin. (c’-e’) Magnification of the hub region. (c, c’) wildtype (*dMef-Gal4* crossed with *w^1118^* flies). (d, d’) *plexA*^RNAi^ (e-e’) *sema1b*^RNAi^. (f-g) Quantification of adult gap-defects. (f) Manually-traced gaps in plexA^RNAi^ as an example (same testis as in (d)). (f’) Quantification of the percentage of uncovered area after different *plexA* and *sema1b* genetic perturbations. (g) Computationally unrolled testis from (f), with a histogram curve displaying grey values of gap-data proportional to gap size. (g’’) These values plotted as a heatmap for multiple replicates for two different *plexA*^RNAi^ and *sema1b*^RNAi^ lines. n=8 for each genotype. (related to Fig S1)

While cell-cell contacts are dynamic during migration, a connected cell sheet must be actively maintained to ensure full muscle-coverage to allow organ-sculpting[11, 15]. Gap closure is supported by N-Cadherin-based cell-cell adhesion, free-edge protrusions and wound closure-like purse-string-dynamics [10]. Remaining gap-free is also crucial during sprouting angiogenesis and vascular repair, during which endothelial cells are surprisingly dynamic and undergo free-edge based collective migration[16–20]. The endothelial monolayer must maintain integrity while migrating. Collectively migrating endothelial cells actively avoid gaps using junctional lamellipodia[5, 21], reminiscent of mesenchymal cell-cell edges in TNM. Given commonalities like these, another key challenge for the field is to define conserved mechanisms by which cell-cell contact-signaling ensures gap-closure during migration.

One advantage of the TNM system is that migration defects can be identified by analyzing adult testis morphology. Only continuous muscle coverage allows normal organ-sculpting (Fig 1b, b’)[15], reminiscent of how airway smooth muscles shape epithelial branches in mammalian lungs[22, 23]. We exploited this to carry out a large-scale candidate screen for myotube-specific regulators of testis shape (BioRxiv). One of the most notable hits was the receptor Plexin A (PlexA). Knockdown led to major gaps in muscle coverage at the distal end of the testis.

Semaphorins and their Plexin receptors are classical axon guidance factors regulating contact-dependent axon-axon repulsion[24, 25] highly reminiscent of CIL[26]. Semaphorin/Plexin signaling is also used in many other processes, including bone formation, immune responses, angiogenesis and vasculogenesis[27, 28]. Clinically, it is implicated in atherosclerosis, diabetes, cancer growth and metastasis[27, 29, 30]. *Drosophila* Plexin/Semaphorin signaling also has roles outside axon guidance. For example, it mediates repulsive cues steering glial cells to migrate on axons in imaginal discs[31], orchestrates collective cell rotation of *Drosophila* follicle cells[32] and allows extrusion of apoptotic cell in wing discs[33].

Plexins are transmembrane receptors with an extracellular Sema-domain and intracellular Rho-binding and Ras/Rap-Gap domains. Their Semaphorin ligands can be transmembrane, secreted, or GPI-anchored, and bind Plexin via their own Sema-domains[28, 34]. Semaphorins on neighboring cells often act as activating ligands, functionally dimerizing Plexins to bring the cytoplasmic domains into proximity. This can activate the Ras/Rap GAP domain, downregulating R-Ras, M-Ras, or Rap GTPases[27, 35]. However, recent structural studies suggest a novel role for *Drosophila* Sema1b. It cannot dimerize[36] but can bind PlexA in cis[37], suggesting Sema1b is a PlexA antagonist rather than an activator[37], but this has yet to be tested in vivo.

We found that PlexinA/Semaphorin signaling plays a key role in allowing cells to avoid gaps during collective cell migration. Our data also provide an in vivo role for PlexA/Sema1B antagonism. PlexA, acting via the *Drosophila* R-Ras homolog Ras2 and the related GTPase Rap2L, maintains cells at the correct place on the mesenchymal-epithelial continuum, facilitating spreading and effective migration under confinement while avoiding gap formation. PlexA and Sema1B antagonism provides this crucial balance.

## Material and Methods

### Fly husbandry

Fly crosses and husbandry were performed according to standard methods[38]. Crosses were performed at 25 °C. Timing of prepupae was performed at 26.5 °C in a cell incubator. All fly stocks used can be found in Supplemental Table 1. We used *w^1118^* for control crosses and referred to it as wildtype or WT. For epistasis-experiments we co-expressed UAS-RFP in single-RNAi controls, to have the same number of UAS-promoters in all conditions, correcting for potential competition for dMef2-Gal4-binding.

### Staining techniques and Imaging

#### Fixed adult testes

We dissected testes from adult flies 1–3 days after hatching in 1.5x PBS and fixed them for 20 mins in 4% PFA in PBS. After three washing steps in 1.5x PBS and one in 1.5x PBS with 0.1% Tween 20, we stained testes overnight in 1:500 Phalloidin (*ThermoFisher, Alexa Fluor 488*) in 1.5x PBS with 0.1% Tween 20 at 4°C. After three washing steps, we imaged them in 35 mm glass bottom imaging dishes in PBS, using a LSM Pascal (Zeiss) with a 10x (Zeiss EC Plan-Neofluar 10x/0.3) dry objective. Pictures shown of adult testes are maximum intensity projections of large z-stacks. We improved the contrast in Fiji to make all structures visible.

#### Immunofluorescence on fixed pupal testes

We staged pupae by collecting prepupae and letting them develop for 30 h at 26.5 °C which leads to a similar coverage as 33 h at 25 °C. We dissected pupal testes from prepupae in 1.5x PBS and fixed them in 4% PFA in PBS. After three washing steps in 1.5x PBS and one in 1.5x PBS with 0.1% Tween 20, we stained testes overnight in 1:500 Phalloidin (*ThermoFisher, Alexa Fluor 568*), primary antibody and DAPI (*ThermoFisher*) in 1.5x PBS with 0.1% Tween 20 at 4°C. We used the following antibodies: anti-N-Cadherin (*DSHB, DN-Ex #8*, 1:500), anti-myc (Cell Signaling Technologies, 71D10, 1:200) and anti-GFP (Abcam, ab290, 1:1000). We used secondary Dylight secondary antibodies (*ThermoFisher*). We mounted testes in Fluoromount-G (*ThermoFisher*) and imaged them using a Zeiss LSM 880 with Airyscan, an inverted stand and a 63x/1.4 oil Plan Apochromat objective.

### Live cell imaging

After timing prepupae and letting them develop 30 h at 26.5 °C, we dissected testes in M3 Medium (Shields and Sang, Sigma Aldrich) with 10% FCS (Thermo Fisher Scientific), and 1x penicillin/streptomycin (Gibco) at room temperature[12]. We mounted them in 0.5% low gelling agarose (Sigma Aldrich) in M3 Medium, and imaged them, using a Nikon Ti2 inverted microscope with Yokogawa CSU-W1 spinning disk and Hamamatsu ORCA-fusion BT sCMOS camera. For over-night live-cell imaging we used a 25x/1.05 Silicone Apochromat objective. For short-term high-resolution imaging we used a 60x/1.4 Oil Plan Apochromat. For over-night images we used Fiji (ImageJ) to change *Gamma* to 0.5, to make all structures visible without oversaturating parts of the tissue. In all other images, we left *Gamma* untouched. Using Photoshop (Adobe) we changed brightness and contrast to make all structures visible.

### Image analysis

#### Adult testes analysis

We used Fiji[39] to manually trace gaps and their total area as “*polygon selection*” ROIs. We used Excel (Microsoft) to measure the quotient of gap area and total area. For the analysis of proximodistal gap-distribution, we turned gap-ROIs into a new channel, with the gaps in white. To define the proximo distal axis, we used the “*segmented line*” tool. We then linearized testes, applying the “*Edit* > *Selection* > *Straighten*” function to this line-ROI with a width of 300 µm and linearized both, the imaging data and the second “gaps-in-black-and-white”-channel. We then applied “*Analyze* > *Plot Profile*” after selecting everything on the “gap” channel, which creates a profile that sums gray values in each vertical line. These values are proportional to the “number” of gaps at each position. We then turned the X-axis values from absolute position values to relative values in Excel and grouped values using the *pivot table* function, to compare multiple testes. We used GraphPad Prism to plot data as a heatmap in which every line corresponds to a single testis.

#### Migration Track analysis

Cell tracks were tracked using Imaris (Andor), by first applying the “*spots*”-module, that after defining the correct parameters, recognized triplets or doublets of nuclei as a single position. Subsequently, gaps and errors by the automatic tool were corrected by hand-tracking. Sometimes, the “*spots*”-module left out single time steps. To do spatial statistical analysis, we had to fill these “gaps”. Therefore, we developed an R-Script, that interpolates missing positions from the “*position*”-output from Imaris. As there is not much movement in a single time-step, interpolation worked very well to approximate the cell position. The script produces a file-format that can be re-entered into Imaris. After using it, we checked that “filled” positions still correctly match the actual cell positions and did all further analysis with these corrected data. To correct for drifting and shaking, we manually applied a “*reference frame*” in Imaris, that stabilizes the testis with its proximal end as a point of reference. For subsequent analysis, we used the “*Position Reference Frame*” output. For motility and angle analyzed we analyzed the timepoints from 0 min – 400 min after beginning of the experiment. To depict cell tracks, we used the online-tool *MotilityLab*[40] after the smoothing process explained later. Same of these methods were published before[11].

#### Motility and Migration Distance analysis

To measure motility, we created an R-script (*R version 4.4.1 (2024-06-14 ucrt), RStudio 2024.04.2 Build 764*) that analyzes the magnitudes of vectors connecting a given cell position (*p*) at each timepoint (*t*) to its position 50 minutes later: 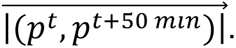 When comparing shorter periods, movements that are independent of migration – like the entire system slightly shaking or movement of the nucleus within the cell – have stronger effects than the actual motility. Therefore, we used this “rolling average”-method. The total displacement is the magnitude of a vector between the first and the timepoint 400 min later: 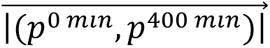 and was measured using Excel.

#### Directionality analysis

To correct for the errors caused by the slight shaking and by nuclear motion, we smoothed the tracks using an algorithm that finds the smoothest trajectory within a 10 µm radius around each cell position. This method decreases weak effects rather than creating differences. Thus, it decreases the chance of false positive “hits” because of effects like shaking during image acquisition. Since the migration occurs on a curved substrate, angle analysis cannot be performed on 3D Data. Therefore, we first “flattened” the position data using a cylindrical Mercator-projection-like algorithm, resulting in 2D trajectories. These analyses were performed in Excel. The Excel sheets and a detailed description how they work can be found in our previous publication[11].

For angle analysis, we developed an R-script that measures the angle between the proximodistal axis (Fig 2a’) and each vector connecting a cell position at timepoint t to the following timepoint: 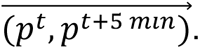 This script uses the ggPlot2 package to plot angle distributions in a rose plot diagram. For backwards motility, we only plotted values from - 160 °C to +160 °C in GraphPad Prism.

**Figure 2.**
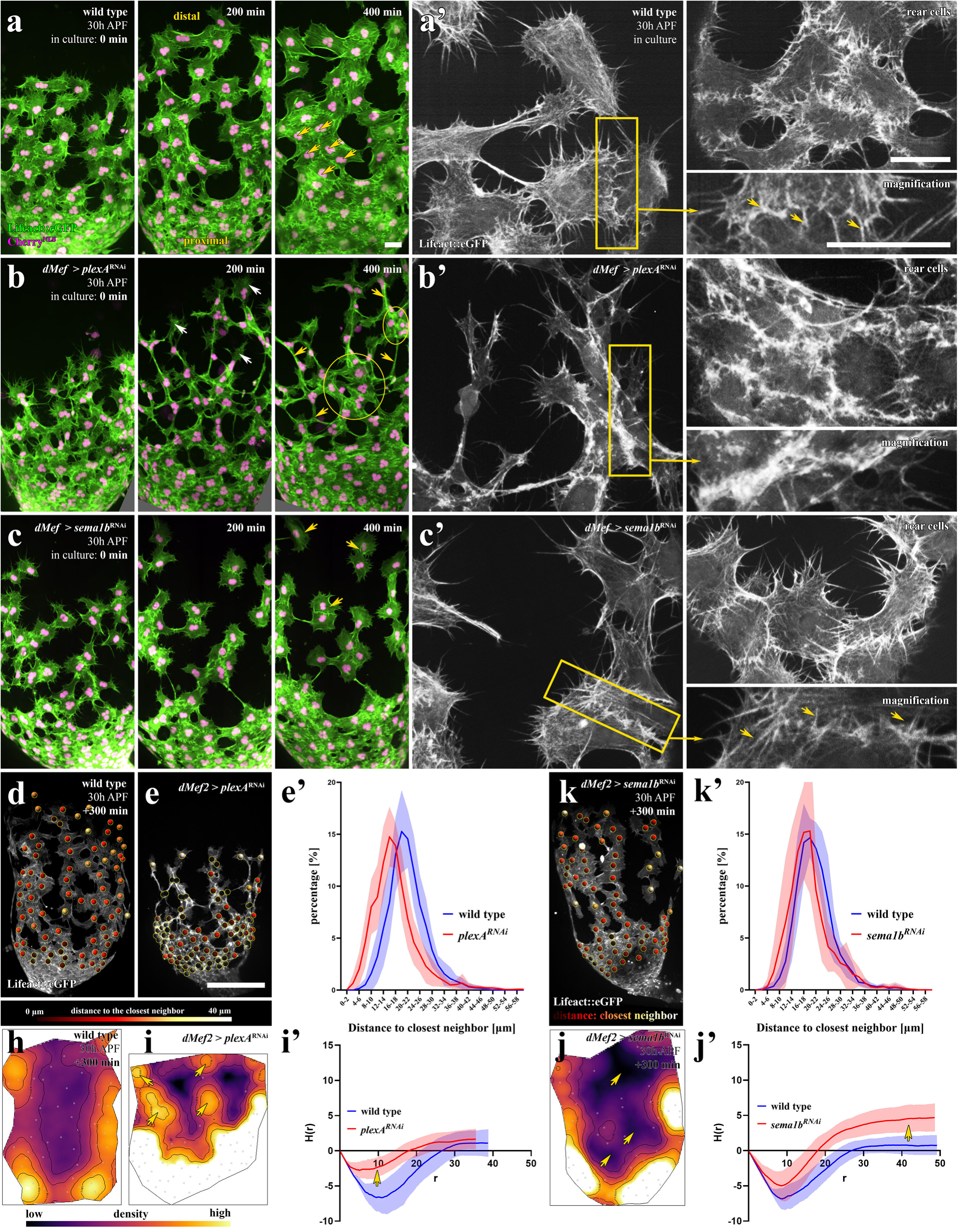
*plexA*^RNAi^ and *sema1b*^RNAi^ both lead to gap formation via different effects on collective cell migration. (a–c) Ex-vivo culture of 30 h testis. TNM express *Lifeact::eGFP* and *Cherry^nls^* under the control of *dMef2-Gal4*. (a) Wildtype, (b) *plexA*^RNAi^, (c) *sema1b*^RNAi^. (a) Yellow arrows: cells with equal cell-cell distances (b): White arrows=growth cone-like structures, yellow arrows=interconnecting cell processes, circles=dense cell clusters. (c) Yellow arrows=dispersed cells. (a’-c’) High magnification live cell imaging, focused on leading cells (left) and more rear cells (left). Lower right: magnification of boxed regions in front cells. Yellow arrows: interdigitating filopodia. Rear cells are from the same videos. Scale bars=20 µm. (d–k) Distance to closest neighbor quantification for wildtype (d), *plexA*^RNAi^ (e) and *sema1b*^RNAi^. Performed at 200–400 min, when quantified cells had left the seminal vesicle. (k). Overlay of micrographs with color-coded dots, indicating the distance to closest neighbor. (e’, k’) Quantification of percentage of cells at given distances to a neighbor. Curve is the mean of 8 curves for 8 testes with ∼ 60–100 quantified cells/testis. Light-colored area = SD. (h-j’) Cell distribution spatial statistics. (h, i, j) Kernel density map showing local cell density of (h) wildtype, (i) *plexA*^RNAi^, yellow arrow=local densifications, (j) *sema1b*^RNAi^, yellow arrows=empty spaces. (i’, j’) Ripley’s H function. (i’) yellow arrow=local densifying effect. (j’) yellow arrow=global homogeneity effect. (detailed explanation in Supp. Fig 3b). Mean of n=8 testes. Light-colored area=range.

#### Nearest Neighbor Analysis

We used Imaris to obtain a list of the nearest neighbor distances at each timepoint. We then used Excel to filter timepoints and used the “*pivot table*” function to generate a histogram-distributions of distances that we transformed from absolute to relative values, to compare replicates. We used GraphPad Prism to plot data and to show the standard deviation as shaded area.

#### Focal Adhesion quantification using Fiji Macro

To quantify integrin adhesion distribution, we developed multiple Fiji Macros in the ImageJ Macro-language. As a first step, integrin adhesions marked in GFP must be hand-clicked using the “*multi-point*” tool and saved in the ROI manager. One of the tools then uses the Fiji “*threshold*” function, to obtain a black and white mask of the cells, to then create a ROI of their “outline”. We used different threshold algorithms, depending on the sample, to find an optimal ROI, that resembles the real outline that is clearly visible. The tool does that for the GFP and the RFP channel and creates a sum-ROI, which experience showed us to give the best results capturing the actual tissue boundaries. Another tool then creates lines between each point of the hand-clicked multipoint and the closest point on the boundaries based on a preexisting macro[41]. This is shown in Figure 6 a/b. The tool gives the length-values of these lines as an output. Another tool creates 10,000 random points within the cell boundaries, that are analyzed the same way as the measured values, to later normalize for tissue geometry. We used the *pivot table* function in Excel to create a histogram and to turn absolute values into relative values to compare multiple conditions. The distributions are shown in Supp. Fig. 5 b and d. We also used Excel to calculate the observed-to-expected ratio explained in Supp. Fig. 5 e and depicted in f.

#### Thickness Quantification

To analyze thickness, we used the Imaris XTension Biofilm Analysis on 60x short-term live cell imaging data (https://imaris.oxinst.com/open/view/biofilm-analysis). We plotted the output in GraphPad Prism.

#### Spatial Statistics

To analyze distributions of cells during long-term imaging and to apply Ripley’s H, L, K-function[42], we created an R-script using the comprehensive spatial statistics package *spatstat*[43] To create an owin-window we used the *alphahull* package[44] to obtain an *alphashape* (a type of concave hull, that closely resembles the actual tissue boundaries). We used a preexisting function to turn this object into a spatstat-readable *owin* object (https://github.com/EricMarcon/SpatDiv/blob/HEAD/R/alphahull.R). The script then applies Ripley’s H-function to each timestep within a defined range (We also implemented K, L, J and pcf-function), then calculates average values, first over time and then over replicates. It also gives the upper and lower boundaries of the total range as an output. We decided to use *Ripley’s Isotropic Edge Correction* (iso) which for simplicity reason, is the only “edge correction” that the script creates as an output table.

To create kernel density maps, we created a second R-script that uses the same *alphashape* algorithm on every timepoint and then uses the density function to create a kernel-density map for each single timepoint, using a *Epanechnikov* (parabolic) kernel, whose bandwidth (*sigma*) is the median of the “*nearest neigbor distances*”, multiplied by 1.2. The script overlays contours in black and cell positions as gray circles. It creates a folder for each replicate and saves a .png for each timepoint. We then used Fiji/ImageJ to create videos from these maps.

#### Statistics

Statistical analysis was carried out in GraphPad Prism. When comparing two conditions, we used Student’s T-Test. When comparing multiple conditions, we used ANOVA. Exact P-values are mentioned in the figure legends. We plotted motility measurements as “*superplots*” to show and analyze averaged replicate values without losing single track values, that would constitute pseudo-replication if analyzed as single measurements[45]. We used GraphPad Prism to perform Student’s T-Test exclusively on the actual replicates. Rose plots (angle distributions) were plotted using the package *ggplot2* in R. Kernel-densities were plotted using standard plot function in R.

#### Detailed author contributions

Maik Bischoff conceived the project, designed the experiments, carried out many of the experiments, and interpreted the results. Jenevieve Norton was trained by Maik and then assisted in most aspects of the project. Maik Bischoff wrote the paper with editorial contributions from the other authors.

## Results

### PlexA signaling in migrating myotubes is important for testis sculpting by ensuring continuous muscle coverage

During a genetic screen we identified *plexA* as a gene whose knockdown affects adult testis morphology (BioRxiv). We hypothesized that this reflected a role for PlexA as a regulator of TNM collective cell migration. We first looked closely at the adult defects. Phalloidin staining reveals that the wildtype testis is completely covered in muscle (Fig. 1c, c’). Muscle-specific *plexA* RNAi-based knockdown causes gaps in the muscle sheet above the distal hub (Fig 1d. d’), reminiscent of previously described defects in migration or adhesion (Fig 1b’)[15]. To quantify testis coverage defects, we manually traced both the total testis area and the area of gaps (Fig 1f), calculating the percentage of the testis not covered by muscles (Fig 1f’). While wildtype testes were always 100% covered, after *plexA*^RNAi^ 5-7% of the testis was uncovered (Fig 1f’). To rule out off-target effects, we tested three UAS-RNAi constructs and two sgRNA-constructs for somatic CRISPR/Cas9 knockout, expressed using the muscle-specific driver *dMef-Gal4* (Fig 1f’)[46]. All yielded the same characteristic defects and phenotypic strength, suggesting knockdown was very efficient and close to a total knockout (Fig 1f’).

TNM migrate from proximal to distal. *plexA*^RNAi^ led to gaps in distal muscle coverage but left the proximal testis covered by muscle (Fig. 1d, red vs yellow arrows). To precisely define where gaps were located along the proximal-distal axis, we computationally unrolled multiple testes and plotted the distribution and size of gaps in a heatmap. This confirmed that after *plexA*^RNAi^ gaps are confined to the distal testis (Fig 1g-g’’).

While PlexA often functions as a receptor, it can also act as a ligand, for example, activating reverse Semaphorin signaling[47]. Therefore, we tested a hypomorphic *plexA* mutant lacking the entire intracellular domain (*plexA*^Δcyto^). *plexA*^Δcyto^ mutants are viable, unlike the null mutant[48]. Homozygous *plexA*^Δcyto^ mutants share phenotypes with our knockdown-strategies, suggesting PlexA acts as a receptor in TNM (Fig S1a,f’). Using endogenously myc-tagged PlexA, we found PlexA is strongly enriched in TNM compared to other testis tissues (Fig S1e–g), reinforcing the idea that PlexA functions in TNM in a tissue-autonomous fashion.

### The effects of *sema1b*^RNAi^ on morphology are similar but not identical to *plexA*^RNAi^

To identify potential PlexA-binding Semaphorins, we knocked them down in all tissues contacting myotubes, revealing that Sema1b has important tissue-autonomous functions (Supplemental Results 1). A recent structural study found that Sema1b–unlike other *Drosophila* Semaphorins–acts as a PlexA-antagonist. However, it remained unknown what roles, if any, PlexA-Semaphorin antagonism plays in vivo. We first compared the effects of muscle-specific *sema1b*^RNAi^ and *plexA*^RNAi^ on adult morphology (Fig 1d,e). Superficially, the adult defects resemble one another, with both causing gaps above the distal hub (Fig1d,e red arrows; d’,e’). However, we observed one striking difference in comparison to *plexA*^RNAi^: *sema1b*^RNAi^ also causes holes in the muscle sheet in the more proximal testis (Fig 1e, yellow arrows), reminiscent of adhesion defects upon *N-Cadherin*^RNAi^[15].

We developed two working models (Fig S2a): 1) Sema1b inhibits PlexA signaling via its classical pathways, while activating PlexA to signal via an alternate pathway that does not require Semaphorin/Plexin dimerization. 2) Sema1b is an antagonist fine-tuning PlexA activity. Reducing Sema1b causes PlexA-overactivation, and both too little and too much Plexin signaling disrupt migration. To distinguish these, we explored the cell biological basis of their effects on migration.

### *plexA*^RNAi^ dramatically alters cell morphology during collective cell migration

To define how PlexA regulates collective TNM behavior and morphology, we employed 4D-spinning disc microscopy to live image TNM during migration (Fig 2a–c, Video S2). Migrating wildtype TNM move from proximal to distal. Cells are well spread on the substrate, moving while being positioned roughly equidistant to all their neighbors, to cover the testis surface evenly (Fig 2a, Video S2). Expressing *Lifeact::GFP* allowed us to examine F-actin dynamics. Wildtype leading cells send out filopodia, but cell bodies do not dramatically elongate, and at cell-cell borders cells are joined by interdigitating filopodia (Fig 2a’, yellow arrows).

*plexA*^RNAi^ dramatically altered TNM shape and arrangement. Cells were no longer evenly distributed but instead clustered in local groups, with apparent reductions in apical cell area in these regions (Fig. 2b, yellow circles, Video S2: yellow circles). These abnormal cell clusters were connected by long cellular processes or extremely stretched cells (Fig. 2b), resulting in a tug-of-war-like behavior. This caused sudden changes of direction as some cells are retracted by neighbors or pulled into dense clusters (Video S2, white arrows). Some leading cells had growth cone-like leading edges that migrated away from their cell bodies (Video S2, red arrows). Gaps in migrating TNM were strongest at the front of the sheet, consistent with the adult phenotype being restricted to the distal hub (Fig 1d,g). While leading cells retained filopodial protrusions, protrusions at cell-cell borders were dramatically altered, with loss of distinct filopodia and intense actin accumulation (Fig 2b’, Video S3).

To quantify changes in cell dispersion versus clustering we used two approaches. First, we examined the distance of each cell (identified by nuclei centroids) to its nearest neighbor and plotted the distribution of these distances. On average, after *plexA*^RNAi^ cells had a reduced distance to their closest neighbors (Fig 2d,e,e’; Video S4). However, this did not distinguish between local or global dispersion. To directly assess clustering, we used spatial statistics. We created kernel density maps for wildtype and *plexA*^RNAi^, revealing the local cell density (Fig 2h,I, Video S5). While wildtype cells are relatively evenly dispersed, after *plexA*^RNAi^ cells form distinct clusters (light areas) and there are more gaps (dark areas). We next employed Ripley’s H function (H(r)) to assess spatial homogeneity (details are in Supp. Fig. 3). This assesses whether there are more or fewer cells in a circle around each cell, relative to what is expected in a random distribution. It then expands the circle, calculating this measure at each radius r (Supp. Fig. 3b). This revealed that after *plexA*^RNAi^ there were more cells than wildtype at small radii, while at longer distances they became very similar to wildtype (Fig 2i’). This reveals local clustering relative to wildtype. Thus *plexA*^RNAi^ dramatically alters cell and tissue morphology and behavior during migration.

### Depleting *plexA* or *sema1b* has very different effects on collective cell behavior

To discriminate between our two hypotheses for Sema1b function, we compared the effects of *plexA* and *sema1B* knockdown on TNM behavior and morphology in detail. *sema1b*^RNAi^ affects TNM cell shape and arrangement drastically differently than *plexA*^RNAi^. *sema1b*^RNAi^ TNM were similar in overall cell shape to wildtype. Cells were well spread on the substrate, and no dense cell clusters were apparent (Fig. 2a vs c; Video S2). *sema1b*^RNAi^ also did not alter the interdigitated filopodia on cell-cell borders (Fig 2c’), in contrast to *plexA*^RNAi^. Instead, *sema1b*^RNAi^ appears to affect cell-cell adhesion. While wildtype cells remain attached to more proximal neighbors, after *sema1b*^RNAi^ individual or groups of cells are often detached from the sheet (Fig 2c, Video S2), with large gaps opening between TNM (Fig 2c, Video S2), consistent with muscle gaps in adult testes not being restricted to the tip (Fig 1e). This resembles the effect of *N-cadherin*^RNAi^[11, 15].

*sema1b*^RNAi^ also differed from *plexA*^RNAi^ in the effect on cell dispersion. The average distance to the closest cell neighbors after *sema1b*^RNAi^ remained unchanged relative to wildtype (Fig. 2d vs k,k’). The large gaps opening during migration were visible in kernel density maps (Fig. 2j, dark areas, Video S4), with no distinct local clusters. Ripley’s H function-results differed from those of *plexA*^RNAi^. Locally, *sema1b*^RNAi^ and wildtype were similar, but they diverged at longer radii, reflecting greater global heterogeneity (Fig. 2j’, details in Fig S3). In conclusion, our data suggest reducing *sema1b* or *plexA* leads to gaps in the adult muscle sheet by different mechanisms. *sema1b*^RNAi^ causes cell-cell detachment while *plexA*^RNAi^ leads to cell clustering, consistent with a model in which Sema1b antagonizes PlexA.

### Overexpressing PlexA affects morphology by causing adhesion defects more closely resembling *sema1b*^RNAi^ rather than *plexA*^RNAi^

If Sema1b is a PlexA-antagonist, its absence should result in PlexA-overactivation. Indeed, overexpressing wildtype PlexA caused morphological defects in adult testes, by disrupting distal muscle coverage (Fig 3a). Interestingly, expressing a construct lacking the intracellular domain, which should not be able to signal downstream, did not yield a phenotype (Fig 3b), even though it can act as a dominant-negative in other contexts[33, 49]. This is consistent with our finding that PlexA’s cytoplasmic domain is important for its function in TNM. Muscle-specific expression of a non-inactivatable PlexA^SA^ [50] led to severe adult testis morphology defects, resulting in complete loss of musculature in the hub (Fig 3c).

**Figure 3.**
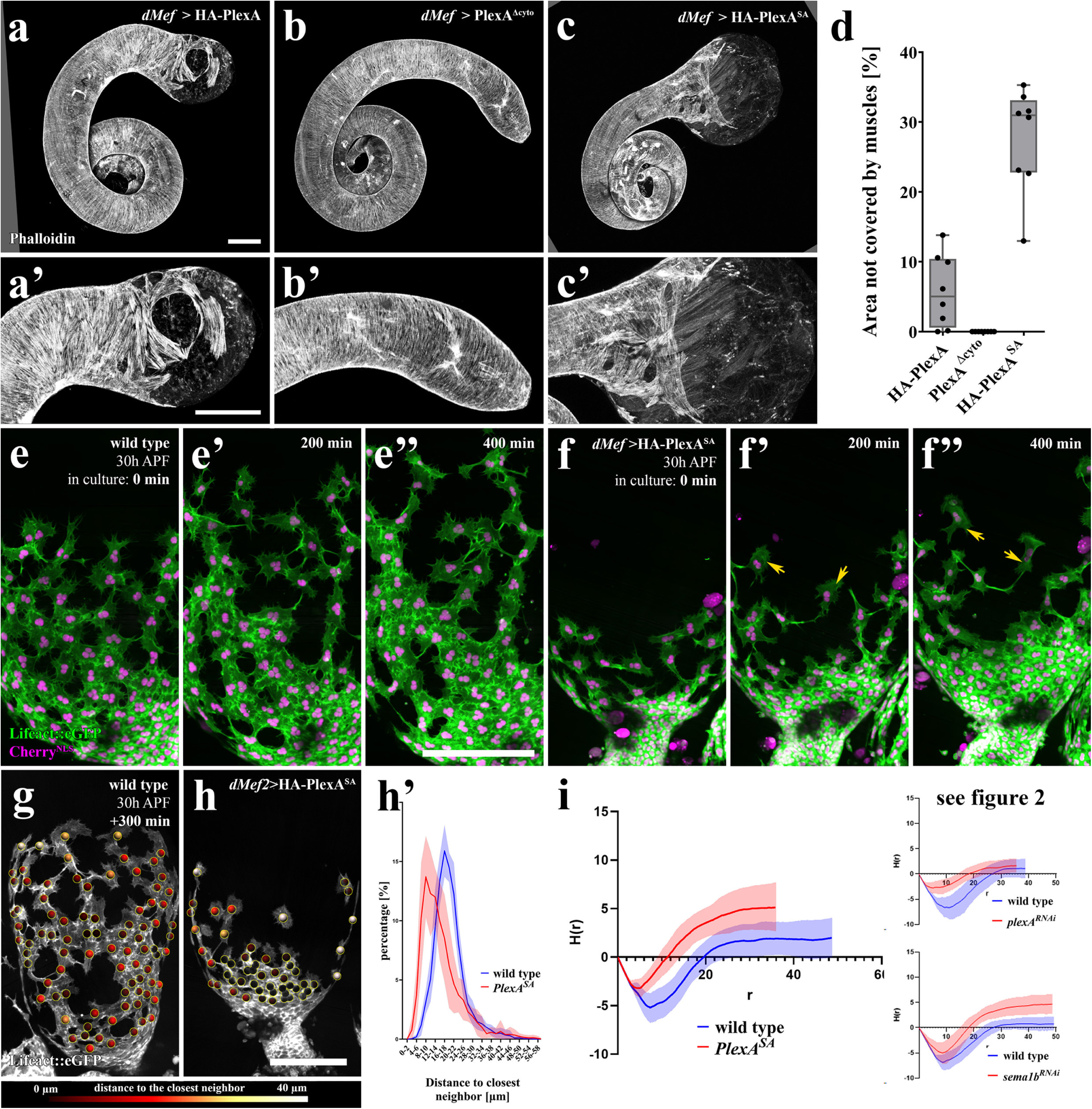
Overexpressing non-inactivatable PlexA causes strong behaviour and cell-cell adhesion defects during collective cell migration. (a–c) Adult testis defects caused by overexpression of wt HA-tagged PlexA (a), PlexA^Δcyto^ (b) and non-inactivatable PlexA^SA^ (c). Scale bars=100 µm. (d) Quantification of uncovered area. Boxplot. Line=Mean and quartiles = box and range (e–f) Ex-vivo culture of 30 h testis. TNM express *Lifeact::eGFP* and *Cherry^nls^* under the control of *dMef2-Gal4*. (e) Wildtype. (f) *PlexA*^SA^ overexpression. Yellow arrows=dispersed cells. (g–h’) Distance to closest neighbor quantification. Performed at 200–400 min, when quantified cells had left the seminal vesicle. (g, h) Overlay of micrographs with color-coded dots, indicating distance to the closest neighbor. Scale bar=100 µm. Quantification in (h’) as a distribution curve of the mean of 8 curves for 8 testes with ∼ 60–100 quantified cells/testis. Light-colored area=SD. (i, i’) Ripley’s H function (detailed explanation in Supp. Fig 3b). Mean of n=8 testes. Light-colored area =range. (right side) Ripley’s H function graphs from figure 2 for direct comparison.

For migration analysis, we used PlexA^SA^, as it caused the strongest adult phenotype. We performed live cell imaging during pupal development (Video S6). Wildtype cells move forward together in well-spread groups, maintaining cell-cell contact with proximal neighbors. In contrast, after PlexA^SA^ overexpression, cell movement was much less collective. Cells could be divided into two groups. Some leading cells detached from the sheet and became isolated, consistent with cell-cell adhesion defects (Video S6, Fig 3f-f’’). This feature was shared with *sema1b*^RNAi^. Many other PlexA^SA^ cells remained at the proximal end of the testis, so the average distance to the closest neighbors is strongly reduced (Fig 3g–h’). Perhaps if leading cells lose adhesion to their proximal neighbors, they cannot pull and spread the sheet effectively. As we will see below, however, the rear cells are still migrating.

When we used Ripley’s H function, wildtype and PlexA^SA^ overexpression TNM diverged at both short and long distances, reflecting greater heterogeneity at all levels after PlexA^SA^ overexpression (Fig 3i). By this measure, the effects closely resemble *sema1b*^RNAi^ (details in Fig. S3), with an earlier divergence due to the dense proximal-population. The effects differ from *plexA*^RNAi^, which converges with wildtype at longer radii (Fig 3i). These data are consistent with Sema1b-PlexA antagonism but suggests expressing PlexA^SA^ causes an even greater PlexA activation than Sema1b^RNAi^.

### *plexA*^RNAi^ causes strong backwards motility while *sema1b*^RNAi^ and PlexA^SA^ do not

To understand how the different alterations in cell clustering in *plexA*^RNAi^, *sema1b*^RNAi^ and PlexA^SA^ cells influence collective cell migration, we compared migration speed and directionality. We tracked cells as they migrated from 0–400 min (Fig 4a). All treatments reduce both migration speed and distance, consistent with the adult phenotypes (Fig 4 b,c,d, details in Supplemental Results 2).

**Figure 4.**
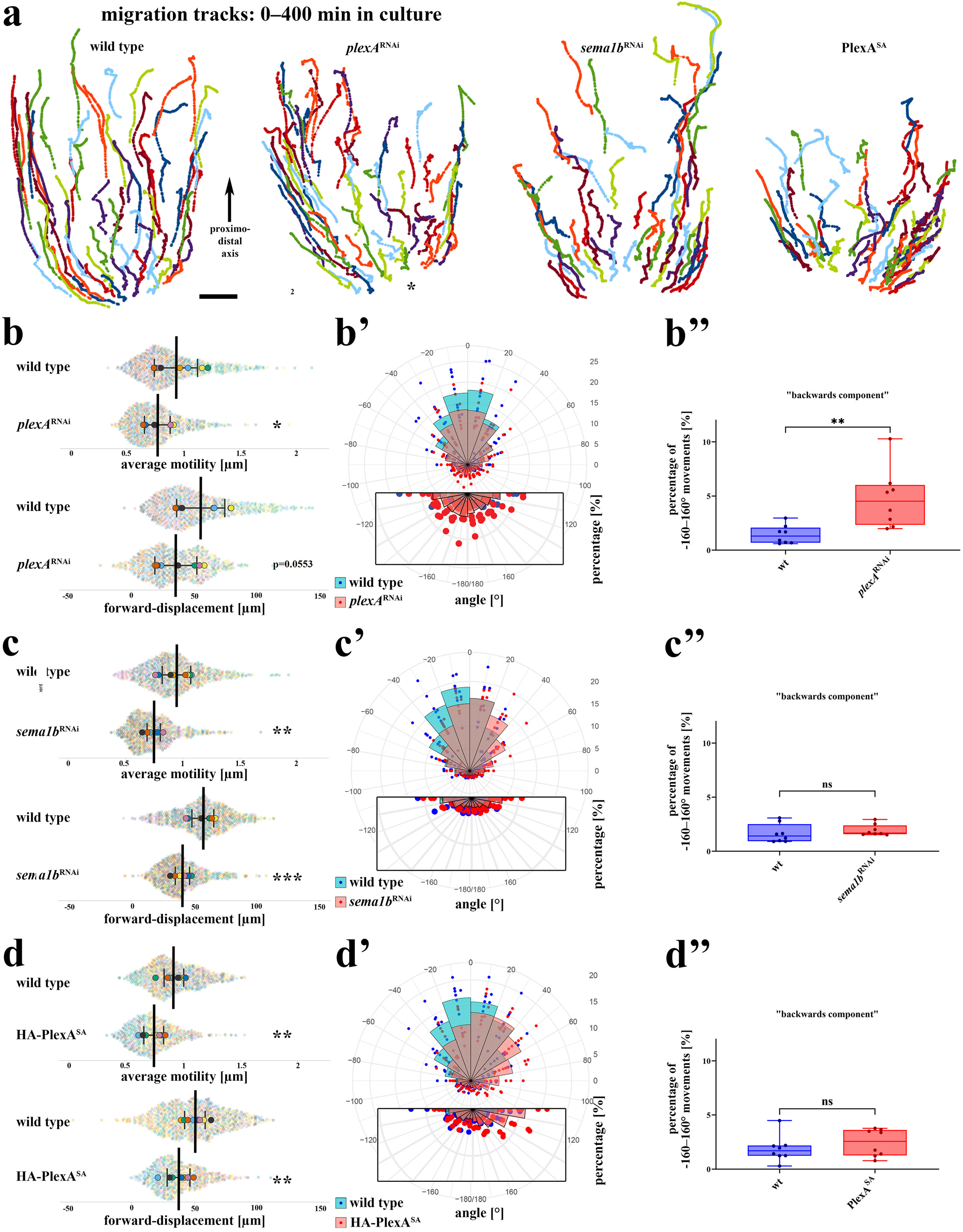
*plexA*^RNAi^, *sema1b*^RNAi^ and PlexA^SA^ overexpression all decrease speed and directionality but only *plexA*^RNAi^ causes reverse migration effects. (a) 4D migration tracks of myotubes in wildtype, *plexA*^RNAi^, *sema1b*^RNAi^ and *PlexA*^SA^ overexpression. Black arrow: proximodistal axis. (b, c, d) Average motility and forward displacement measurements as super plots. n=8 testes (each a big dot) with ∼ 60–100 quantified tracks/testis (small dots). Lines are mean and 95%CI. Statistical tests: unpaired student t test. P values: (b, top): 0.0423, (b, bottom): 0.0553, (c, top): 0.001, (c, bottom): 0.0006. (d, top): 0.0011 (d, bottom): 0.0093. (b’, c’, d’) Rose plot of angle of migration relative to the proximodistal axis during migration (d’). n=8 testes/condition, with ∼ 60–100 quantified tracks/testis at 80 time-steps (400 min). Wedges/Bars=mean of 8 testes, dots: single testes. Bottom: Magnification of the rear-portion of the angle diagram. (b’’, c’’, d’’) Percentage of biased angles between -160°/160°. n=8 testes/condition, with ∼ 60–100 quantified tracks/testis at 80 time-steps (400 min). Student’s T-test.

We next explored whether the different effects on cell clustering had specific effects on cell directionality. Wildtype cells migrate with a strong directional bias along the proximal to distal axis (motion vector angle to the proximodistal axis; arrow in Fig 4a). After *plexA*^RNAi^, *sema1b*^RNAi^ and PlexA^SA^ expression, TNM migrate less directionally (Fig. 4b’-d’). However, the different perturbations differentially affected “backwards” motility (boxes in b’-d’). In wildtype, less than ∼1.4 percent of motion angles are between -160 and 160°. In *plexA*^RNAi^, reversed motion is strongly increased, to ∼4.7 % (Fig 4b’’). This is consistent with *plexA*^RNAi^ cells being retracted into local cell-clusters, causing a characteristic back-and-forth motion (Video S2). In contrast, after *sema1b*^RNAi^ and PlexA^SA^ reversed motion is not significantly changed from wildtype, consistent with our finding that such clusters are absent from *sema1b*^RNAi^ and PlexA^SA^ (Fig 4c’’,d’’).

### PlexA regulates N-cadherin-based cell-cell contacts

We considered the hypothesis that gaps in the migrating sheet and in the final tissue result from altering adhesive properties. We noted above the loss of interdigitating filopodia after *plexA*^RNAi^. To further investigate this, we employed Airyscan2 enhanced resolution microscopy to analyze N-Cadherin-localization at cell-cell junctions. Wildtype cells are joined by junctions in which N-Cadherin localizes along interdigitating filopodia[11], Fig 5a–a’’’’). In contrast, after *plexA*^RNAi^ N-Cadherin foci are not organized along filopodia but instead are linearly distributed along the TNM-TNM border (Fig 5b–b’’’’), thus resembling epithelial cell-cell adherens junctions[51]. We also examined the elongated cellular processes between leading cells upon *plexA*^RNAi^. N-Cadherin staining revealed these are processes of both cells connected via linear punctate junctions along their entire length, reminiscent of axon bundles (Fig 5c–c’’’’’). Thus. PlexA regulates the nature of TNM-TNM cell-cell adhesion.

**Figure 5.**
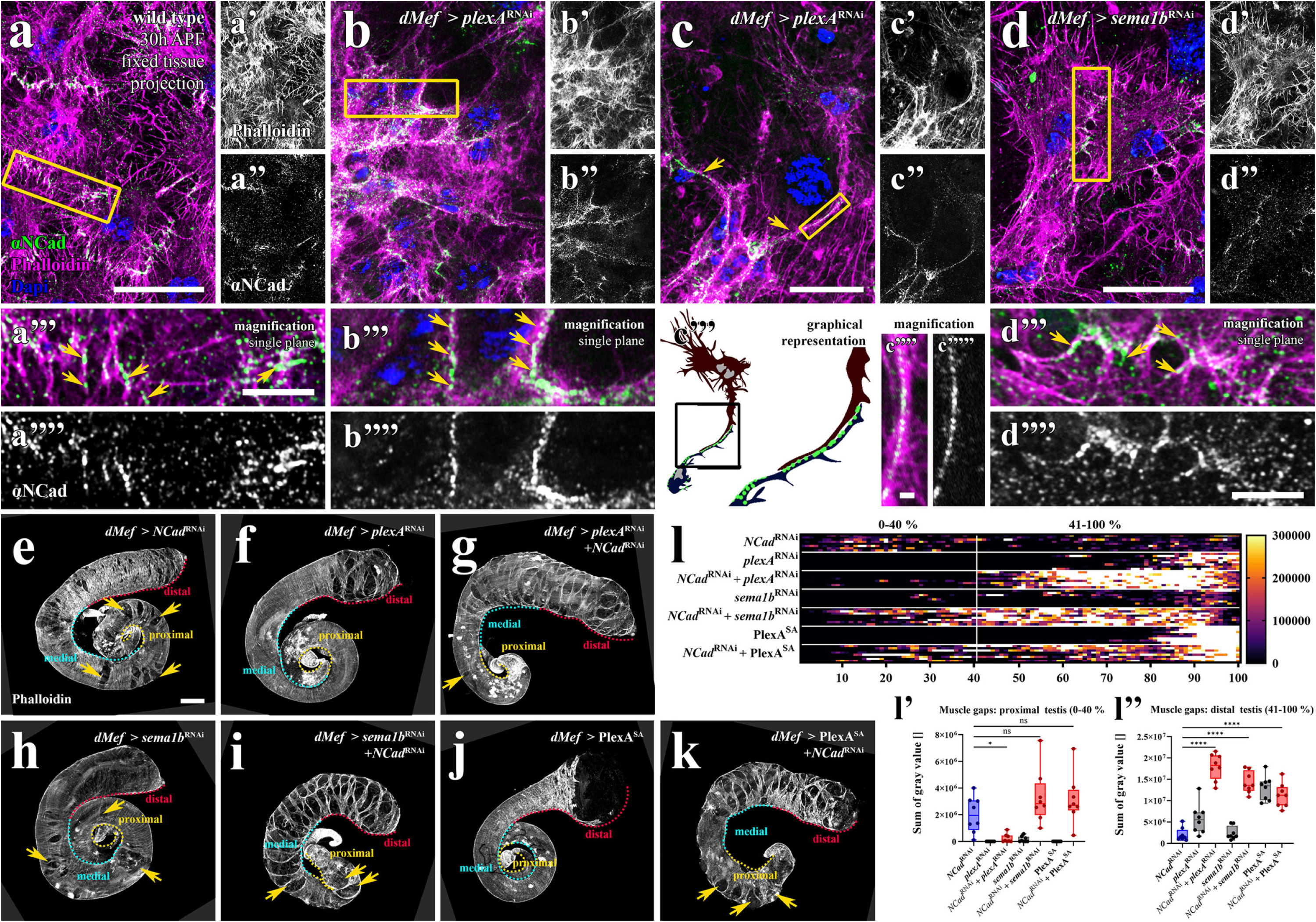
PlexA influences N-Cadherin-based cell-cell contacts whereas Sema1b’s influence on gap-closure seems independent of N-Cadherin. (a-d) Airyscan 2 micrographs of fixed/mounted testis (30 h APF) stained with Phalloidin (magenta), Dapi (blue) and N-Cadherin antibody (green). (a–a’’’’) Wildtype. (b–b’’’’, c– c’’’’’) *plexA*^RNAi^. (d–d’’’’) *sema1b*^RNAi^. Magnifications below represent the yellow boxes above. Axon bundle-like structures in *plexA*^RNAi^ are depicted in (c-c’’’’’) with a graphical representation in (c’’’). Yellow arrows=N-Cadherin-based cell-cell contacts. (e–l’’) Genetic interaction experiments with *plexA^RNAi^*, *sema1b*^RNAi^, PlexA^SA^ and *NCad^RNAi^*. (e–k) Adult testis with different morphological defects. Musculature stained with Phalloidin. Distal, medial and proximal portions of the testis marked. Yellow arrows=gaps in the muscle sheet. Scale bar=100 µm. (l) Gap quantification after computational unrolling (compare figure 1f–g’’). (l’) Sum of gaps in the proximal region (0–40%) (l”) Sum of gaps in the distal region (41–100%)’. n=8 testes/condition. Statistical test: Ordinary one-way ANOVA, Šídák’s multiple comparison test. P-Values: (l’, from top to bottom): 0.1739, 0.0877, 0.0104. (l’’, all values): <0.0001. (related to Fig. S3)

### *sema1b*^RNAi^ and *plexA*^RNAi^ alter cell-cell adhesion in different ways

In contrast to the effects of *plexA*^RNAi^, *sema1b*^RNAi^ had no effect on TNM cell-cell junctions: interdigitating filipodia remained decorated by N-cadherin (Fig. 5d–d’’’’). However, we noted above that *sema1b*^RNAi^ causes a migration phenotype reminiscent of N-cadherin loss[11], even though N-Cadherin is still present. To test if the effects *plexA*^RNAi^ and *sema1b*^RNAi^ on cell-cell adhesion are contrary, we analyzed double knockdown with *N-cadherin*^RNAi^. *N-cadherin*^RNAi^ alone causes strong morphological defects in adult testis morphology with gaps all along the muscle sheet from proximal to distal (Fig 5e)[11, 15]. *plexA*^RNAi^ alone leads to gaps restricted to the distal hub region (Fig 5f). When we examined *plexA*^RNAi^ *N-cadherin*^RNAi^ animals, we saw something interesting: double knockdown substantially increased the size of holes in the distal hub, the region affected by *plexA* single knockdown (Fig 5e–g). Enhanced defects were also observed in the medial region (41–100% of testis length; Fig 5f, l, l’; Fig S4). This synergistic effect suggests PlexA may act in parallel with N-cadherin via different pathways. Interestingly, holes in the proximal testes (0-40 %, testis length) seen after in *N-cadherin* single knockdown were strongly suppressed by double knockdown (Fig 5g,l,l’). Thus, the proximal defects caused by N-Cadherin^RNAi^ are partially rescued by reducing PlexA. Perhaps reducing *plexA* allows the more densely-packed rearward TNM seen after *plexA*^RNAi^ to cope with reduced *N-cadherin* during migration, possibly by stabilizing cell-cell adhesion, consistent with the altered adhesion morphology. *sema1b*/*N-cadherin* double knockdown had quite different effects. It caused synergistic effects *all along* the proximal-distal axis (Fig 5h vs. l,l’’, Fig S4), consistent with the idea that *sema1b*^RNAi^ decreases cell-cell adhesion in a N-Cadherin-independent way. Overexpressing the non-inactivatable PlexA in a *N-cadherin*^RNAi^ background mimics the effects of *sema1b*^RNAi^, with strongly enhanced gaps distally and slightly enhanced gaps proximally (Fig 5k,l-l’’, Fig S3). In summary, PlexA and Sema1b have different and potentially opposite effects on cell-cell adhesion dynamics.

### PlexA is crucial for cells to maintain a flat mesenchymal cell shape during migration

TNM are tethered to the substrate via Integrin-dependent matrix adhesions assembled underlying both free-edge filopodia and cell-cell edge filopodia. Adhesions in free-edge filopodia have a much longer lifetime[11], stimulating cells to extend into the open space and fill it. However, interdigitating filopodia at cell-cell borders also have underlying matrix adhesions, even though they are not extending[11] (Fig 6a,a’’’’ yellow arrows; cell-matrix adhesions were visualized by expressing the focal-adhesion-targeting-domain of Fak fused to GFP in TNM[52]). Strikingly, the number of matrix adhesions at cell-cell contacts appeared strongly reduced after *plexA*^RNAi^ (Fig 6b,b’’’’). We quantified the distance of each individual matrix adhesion from the cell-sheet edge (Fig. 6a’’,b’’, Cartoon in Fig S4a) and plotted the distribution of distances (Fig S5b). Changes in cell clustering caused by *plexA*^RNAi^ mean that even random points are, on average, further from a free edge (Fig S5c). To normalize for this, we created a set of random points (Fig. 6a’’’,b’’’) and analyzed their distance distribution to reveal what a random distribution looks like in each architecture (Fig. S5d). We then used this to calculate whether there are more adhesions or fewer adhesions at cell-cell borders versus those at free edges (Fig S5e). *plexA*^RNAi^ had little effect on matrix adhesion numbers near free edges (distance 0-10µm) (Fig. 6C, top). However, *plexA*^RNAi^ led to a strong reduction in matrix adhesions further from free edges (>10 µm), thus including matrix adhesions at cell-cell borders (Fig. 6c bottom).

**Figure 6:**
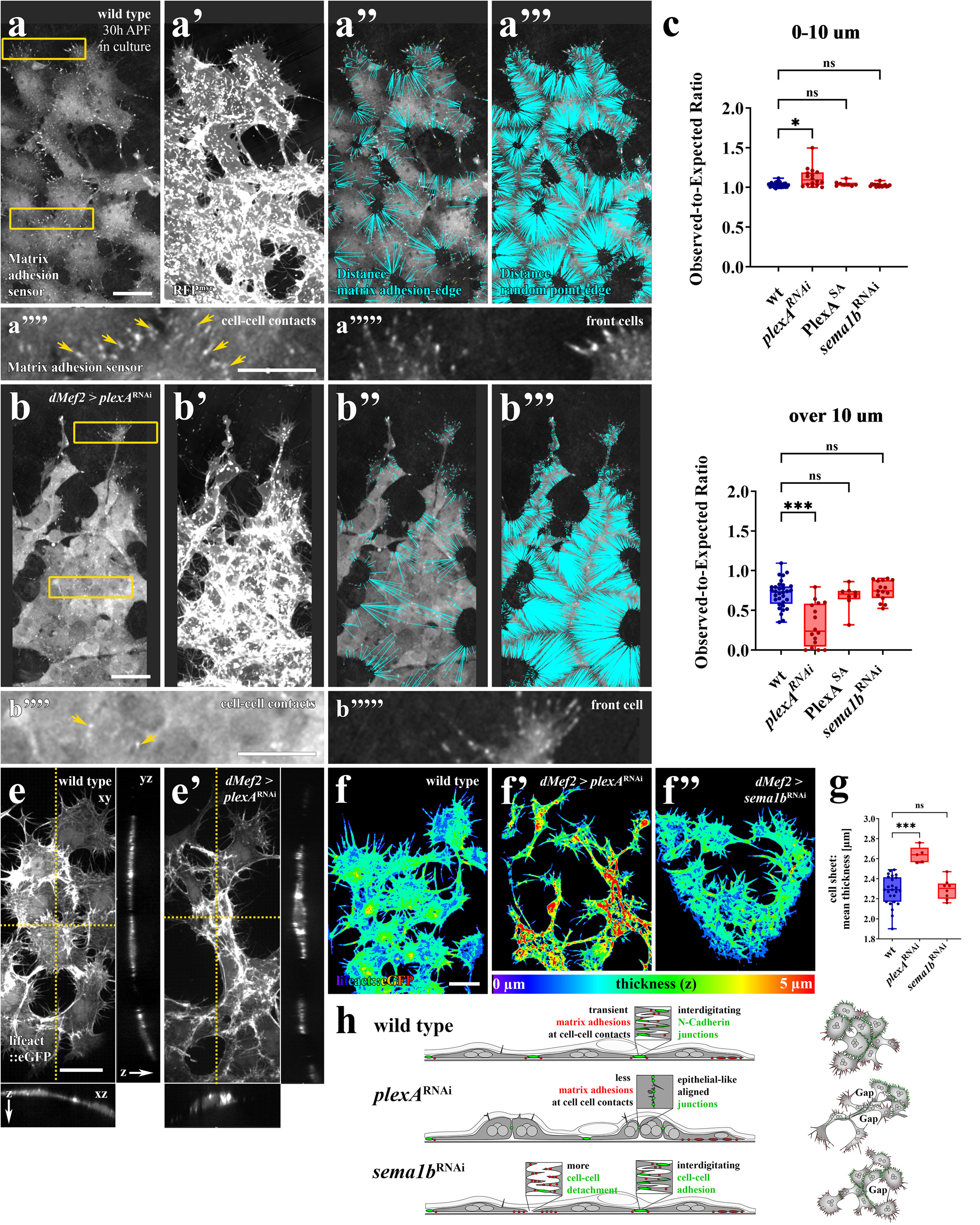
*plexA*^RNAi^ perturbs cell-cell contact adjacent matrix adhesions and alters TNM cell height while *plexA* overexpression or *sema1b*^RNAi^ do not. (a–b) High resolution live-cell imaging of ex-vivo cultured (30 h APF) testis for quantification of Integrin adhesion distribution in wildtype (a-a””’) and *plexA*^RNAi^ (b-b””’) TNM express *Focal adhesion targeting domain (fat)::eGFP* under control of *dMef2-Gal4*. Details of the method are in Fig S4. (a, b) Overview of matrix adhesions. (a‘, b’) myr-RFP revealing cell edges. (a”, b”) Lines displaying distances from matrix adhesions to the nearest cell edge. (a”’, b”’) Lines displaying distances from random points to the nearest cell edge. (a””. a””’, b””, b””’) Closeups of yellow boxes in (a, b) showing either cell-cell contacts or front cells. Yellow arrows: Matrix adhesions near cell-cell contacts. Scale bar (overview) = 20 µm, (magnification): 5µm. Statistical test: Mixed effects-analysis, Šídák’s multiple comparison test. P-Values: (from top to bottom): 0.6454, 0.8397, 0.0201, 0.9750, 0.8481, 0.0003. (related to Fig. S4) (e-g) Quantification of cell height. Wildtype (e, f), *plexA*^RNAi^ (e’, f’) and *sema1b*^RNAi^ (f”). (e. e’) xz and yz sections through a 3D micrographs. Scale bar: 20 µm. (f-f”). Sheet thickness quantified with the Imaris XTension Biofilm Analysis. (g) Mean height of the cell sheet. n= wt: 25, *plexA*^RNAi^: 5, *sema1b*^RNAi^: 8. Statistical test: Kruskal-Wallis, Dunns’s multiple comparison test. Adjusted P-Values: (from top to bottom): > 0.9999, 0.0008. (h) Schematic of cell architecture defects after *plexA*^RNAi^ and *sema1b*^RNAi^. Different effects lead to similar gap formation during collective cell migration.

We next examined cell-matrix adhesions after *sema1b*^RNAi^. In contrast to *plexA*^RNAi^, there was no reduction of cell-matrix adhesions adjacent to cell-cell borders (Fig S5g–j; Fig 6c). We also assessed cell-matrix adhesions after expressing PlexA^SA^. Because of strong cell dispersion due to apparent cell-cell adhesion defects, many *dMef2* > PlexA^SA^ specimens did not have any areas more than 10 µm away from the next cell sheet-edge within the ROIs we analyzed (Fig S4h), and thus only 8/16 specimens could be analyzed. These had a normal distribution of remote matrix adhesions, like *sema1b*^RNAi^ (Fig S5g-j, Fig 6c). Thus, *plexA*^RNAi^ and *sema1b*^RNAi^ have different effects on cell-matrix adhesions.

TNMs migrate through a confined space between pigment cells and cyst cells and thus must maintain a flattened morphology. One role cell-matrix adhesion close to cell-cell contacts might have is to keep the cell flat by dynamically tethering it to the substrate on all edges in a tent stake-like fashion. To test if TNM height is affected by PlexA signaling, we measured cell height after *plexA*^RNAi^. Strikingly, *plexA*^RNAi^ TNM are, on average, much taller, especially within clusters—this is apparent in cross-sections (Fig 6e) and is even clearer when cell height is quantified across the sheet (Fig 6g). This change in cell height is not seen after *sema1b*^RNAi^ (Fig 6f’’,g).

Together, the changes in the cellular architecture upon *plexA*^RNAi^ are consistent with the cell clustering behavior observed during collective cell migration (Fig 6h). These data suggest PlexA acts to keep the cell-cell interfaces more mesenchymal, which in turn keeps cells flat, while still maintaining contact via dynamic protrusions. This facilitates migration under confinement. *plexA* reduction causes cell junctions to become “epithelialized”, with fewer interdigitated protrusions and more linear cell-cell junctions. Cells become taller, also a more epithelial character. Consistent Sema1b acting as an antagonist rather than an activating ligand, *sema1b*^RNAi^ does not phenocopy these “epithelializing” effects (Fig 6h). Instead, *sema1b*^RNAi^ has effects resembling the destabilization of cell-cell adhesion, resulting in gap-formation like that seen after *N-Cadherin*^RNAi^. Further analyses suggest the defects observed upon *plexA*^RNAi^ and *sema1b*^RNAi^ are independent of Myosin activity (Supplemental Results 3).

### Genetic interactions suggest PlexA and Sema1b are in the same pathway

Our analyses revealed that PlexA and Sema1b have opposing effects. This is consistent with them working in two independent pathways or by antagonism within the same pathway. To test which is more likely, we performed epistasis analysis. First, we analyzed muscle-specific double RNAi of *sema1b* and *plexA*. We expected that if they operated via separate mechanisms, we would see synergistic effects on muscle coverage, as we saw with double knockdown of PlexA and Myosin or N-cadherin (Fig 5g; S6f). However, *sema1b*^RNAi^ did not enhance the phenotype of *plexA*^RNAi^ (Fig 7a–d’,e). When we looked at the distribution of gaps along the testis, things became even more interesting. The holes in the medial region of the testis observed upon *sema1b*^RNAi^ (Fig. 7c, arrows) disappeared after *sema1b*/*plexA* double RNAi (Fig 7f,f’) reminiscent of the gap suppression we observed after *plexA N-cadherin* double RNAi (Fig 5l-l’’). This partial rescue of the *sema1b*^RNAi^-phenotype by reducing *plexA* supports an antagonistic relationship. However, there are remaining distal gaps after double knockdown. This may indicate that a precise level of PlexA signaling is required for wildtype cell behavior, and we have not hit this perfectly after double knockdown. Alternately Sema1b might both antagonize canonical PlexA, but also stimulate PlexA signaling via an alternate pathway, or Sema1B might have additional Plexin-independent functions.

**Figure 7.**
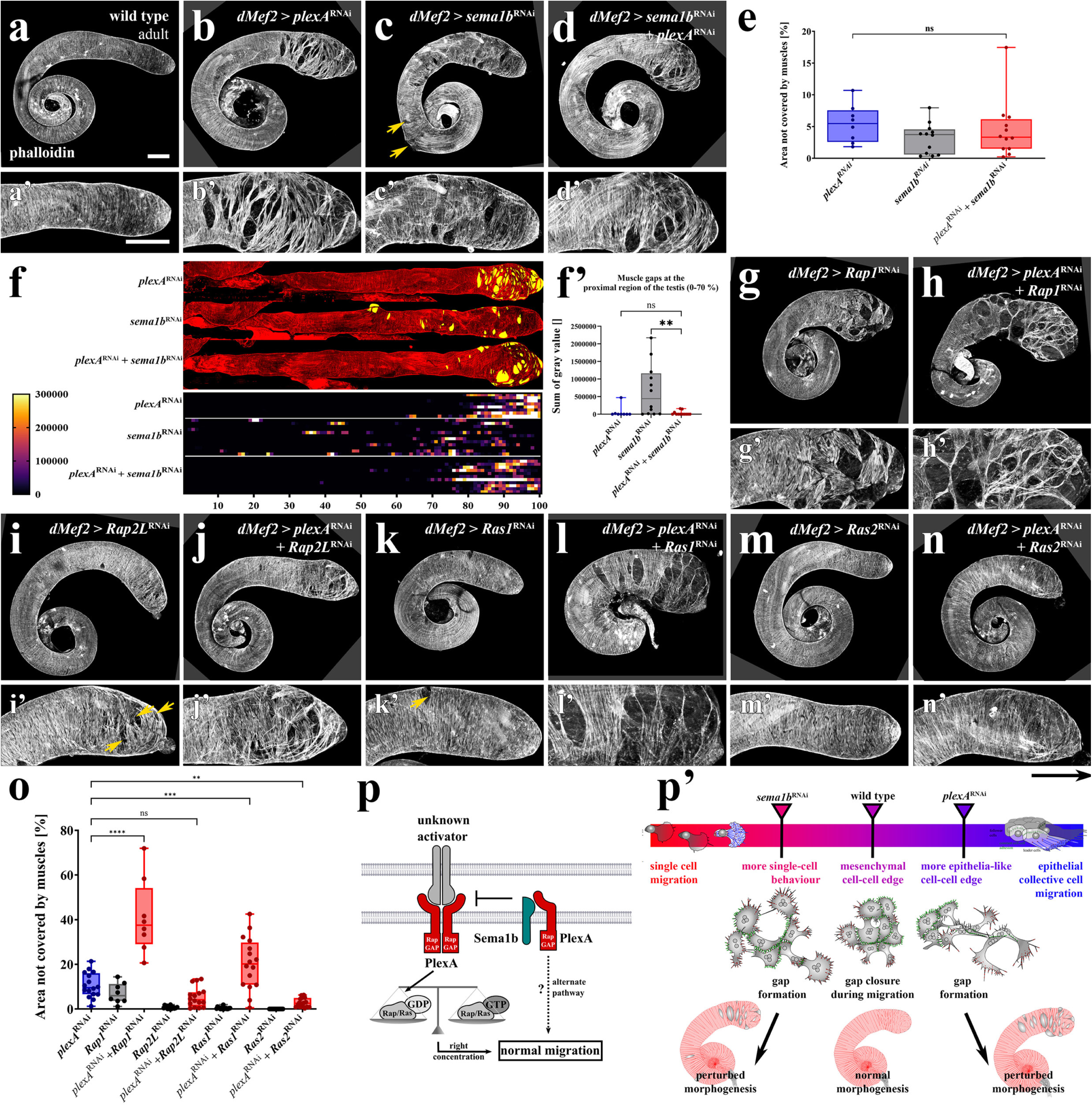
Genetic interactions suggest PlexA and Sema1B are antagonists and PlexA acts as a Rap2L/Ras2-GAP. (a–f’) Genetic interaction experiments with *plexA^RNAi^* and *sema1b*^RNAi^. (a-d) Adult testis with different morphological defects. Musculature stained with Phalloidin. (a’–d’) Magnifications of the hub regions. Yellow arrows: medial gaps. Scale bar: 100 µm. (e) Quantification of area uncovered. *plexA*^RNAi^: n=8, sema1bRNAi, *plexA*^RNAi^ + *sema1b*^RNAi^: n=12. Statistical test: Ordinary one-way ANOVA, Šídák’s multiple comparison test. P-Value = 0.5562. (f) Gap quantification after computational unrolling. (f’) Muscle gaps in the proximal region (0–70%). Statistical test: Ordinary one-way ANOVA, Šídák’s multiple comparison test. P-Values: (from top to bottom): 0.9861, 0.0044. (g–o) Genetic interaction experiments with *plexA*^RNAi^, *Rap1*^RNAi^, *Rap2L*^RNAi^, *Ras1*^RNAi^ and *Ras2*^RNAi^. (g-n) Adult testis with different morphological defects in. (g’–n’) Magnifications of the hub regions. (o) Quantification of area uncovered. *Rap1*^RNAi^, *plexA*^RNAi^ + *Rap1*^RNAi^: n=8, others: n=16. Statistical test: Ordinary one-way ANOVA, Šídák’s multiple comparison test. P-Values: (from top to bottom): 0.0037, 0.0002, 0.0828, <0.0001. (p-p’) Schematic of a proposed PlexA/Sema1b pathway (p) and illustrations of how signaling, cell-architecture, and collective cell migration are altered upon knockdown of *plexA* and *sema1b*.

### Rap/Ras GTPase proteins play roles in migration and some appear to act downstream of PlexA

One prominent mode of PlexA signaling is to act as a GTPase activating protein (GAP) for Rap/Ras-family GTPases upon Semaphorin activation, thus turning down GTPase signaling[27, 53]. Therefore, we first tested if any of the four *Drosophila* Rap/Ras superfamily GTPases[54–56] have roles in TNM migration. *Rap1*^RNAi^ causes defects in distal muscle coverage (Fig 7g,g’), a phenotype roughly resembling that of *plexA*^RNAi^ (Fig 7b,c). *Rap2L*^RNAi^ causes a similar but much weaker phenotype (7i,i’) *Ras1*^RNAi^ causes general defects in shaping, with the testis shorter and broader, and leads to a small number of gaps (Fig 7k,k’), while Ras2^RNAi^ does not cause any defects (Fig 7m,m’).

We examined double knockdown to explore whether these GTPases act downstream of or in parallel to PlexA. *plexA*/*Rap1* double knockdown surprisingly caused strong synergistic effects, suggesting they affect migration via parallel pathways (Fig 7h,h’,o). *plexA*/*Ras1* double knockdown was also somewhat synergistic (Fig. 7l,l’,o), and thus is most consistent with action in parallel pathways. In contrast, *plexA*/*Rap2L* double knockdown did not change the *plexA* phenotype (Fig 7k,k’,o), while *plexA*/*Ras2* double knockdown leads to significant rescue of defects caused by *plexA*^RNAi^ (Fig 7n,n’,o). These data are consistent with the idea that in TNM PlexA acts as a Rap/Ras GTPase for Ras2 and potentially also Rap2L. Consistent with this, co-immunoprecipitation suggested Ras2-GTP binds PlexA much more effectively than Ras1-GTP[50]. Ras2 is a vertebrate R-Ras ortholog[50], a known Plexin target. In summary, these data support the notion that multiple Ras/Rap GTPases act in TNM migration, with gap-closure defects arising from too much or too little Ras/Rap activation.

## Discussion

Epithelial-to-mesenchymal transitions play multiple key roles in development and homeostasis and are altered in many disease states. Here we describe our discovery that Plexin-Semaphorin signaling plays an unexpected role in TNM migration, fine tuning cells on the mesenchymal-to-epithelial continuum to facilitate migration and avoid gap formation.

### PlexA-dependent contact-mesenchymalization is essential for gap-avoidance and organ-sculpting

The goal of TNM migration is to fully enclose and later “sculpt” the testis, so gaps during migration must be constantly closed by N-cadherin-mediated adhesion, actomyosin-based purse-string constriction[10, 11]. Here we describe another form of gap-avoidance regulated by PlexA, which actively maintains mesenchymal cell-cell junctions and adjacent integrin adhesions, keeping cells dynamically spread and distanced during migration, enabling subsequent organ-sculpting. PlexA reduction results in more linear junctions reminiscent of epithelial contacts with reduced ECM-tethering and cell height increases. This leads to local densification, resulting in decreased speed, directionality, and basal area. The resulting gaps perturb sculpting-morphogenesis. Both “global” EMT and MET induction are known roles of different vertebrate Plexin-activating Semaphorins in epithelia[57]. Semaphorin 7a, 4d and 3e can downregulate cell-cell adhesion and motility[58–60]. Consistent with our findings, *Drosophila* PlexA makes E-Cadherin junctions more dynamic in wing disc epithelia, facilitating neighbor exchange, extrusion of apoptotic cells, and wound closure[33].

### A role for Semaphorin1B-PlexA-antagonism in vivo

Semaphorins are generally viewed as activating ligands for Plexin receptors. However, in TNM *Sema1b^RNAi^* has effects different than those caused by the loss of PlexA, as gaps in muscle coverage arise by opposing mechanisms. Our data suggest *sema1b*^RNAi^ makes cell-cell contacts more dynamic, consistent with a more mesenchymal architecture. Thus, cells detach from their neighbors, creating gaps differently. Consistently, two structural studies revealed that *Drosophila* Sema1b cannot dimerize and acts in vitro as a PlexA antagonist[36, 37]. Vertebrate Sema6a uses a similar molecular mechanism to antagonize Plexin in cis, revealing this type of modulation to be evolutionarily conserved. Our data provide the first in-vivo evidence for PlexA/Sema1b-antagonism. However, since *plexA/sema1b* double-knockdown does not fully suppress migration defects, things are likely more complex. Sema1b might also “activate” PlexA signaling via a non-canonical pathway, or reverse signaling by Semaphorins might be involved—these will be important avenues for future exploration.

### In TNM migration Semaphorin/Plexin signaling does not act by repulsion, but still orchestrates emergent cell-behavior

Plexin signaling prominently induces axonal contact-repulsion by deactivating Integrins, causing growth cone-collapse[50, 61], reminiscent of contact-inhibition of locomotion[3, 62]. In *Drosophila* egg chambers, PlexA represses protrusions during follicle cell rotation[32]. In contrast, in TNMs Plexin signaling may instead stabilize Integrins. Perhaps this involves Cadherin-Integrin antagonism[63]. However, there are several examples in which Plexins stabilize Integrins. *Drosophila* PlexB has this role in neuronal dendrites to enable tiling[64], and integrins are stabilized by PlexinB2 in human embryonic stem cells and by Plexin D1 in endothelial cells[65, 66]. PlexA’s role in TNM gap avoidance provides an example of self-regulation of migration distinct from contact-inhibition of locomotion: PlexA signaling causes contact-mesenchymalization and thereby orchestrates another characteristic emergent cell-sheet-behavior.

### TNM and blood vessel repair gap-avoidance mechanisms provide intriguing parallels

We identified the *Drosophila* R-Ras homolog Ras2 as a potential downstream mediator of PlexA signaling during TNM migration. This is intriguing, as R-Ras is a known target of Plexin Gap activity[67, 68] and a master regulator of blood vessel maturation, maintaining barrier function and preventing blood vessel permeability[69]. R-Ras is crucial for EC maturation by stabilizing cell-cell adhesions[70], partly by suppressing VE-Cadherin internalization[71]. These findings are consistent with our observation that *plexA*^RNAi^– potentially increasing Rap2L/Ras2 activity–causes cells to adopt a more epithelia-like morphology. However, in tumor blood vessels, R-Ras stabilization of cell adhesion prevents gap-formation. In TNM, epithelial densification may lead to gaps, as they must remain very flat to cover the testis-surface. Intriguingly, however, R-Ras GAP mutant mice, which effectively have more R-Ras activity, die of internal bleeding with altered endothelial architecture[72]. Thus, even in blood vessels, hyper-stable junctions might lead to gap formation like in TNM.

Semaphorins are important vascular permeability regulators[57] and a direct and fascinating role for Plexin was discovered recently. PlexD1 acts as a mechano-sensor to stabilize Integrin adhesions and maintain vessels upon mechanical stress[66], consistent with our finding that PlexA may maintain matrix adhesions at cell-cell borders. Endothelial cells also remain somewhat mesenchymal at cell-cell contacts, with intermittent lamellipodia associated with VE-cadherin-based junctions[5, 21], and filopodia at some cell-cell contacts[4, 73]. The prevention of full “epithelialization” of cell-cell junctions may be important to avoid gaps in migrating monolayers. In both, TNM and ECs, this might allow cells to react quickly with preexisting protrusions to constantly forming micro-gaps generated by stresses caused by sheet locomotion. Future studies will have to explore if Plexin signaling and downstream inhibition of the R-Ras-axis, are evolutionary conserved regulators of contact-mesenchymalization.

## Acknowledgements

We are grateful to Alexander Kolodkin, Jon Terman, Liquin Luo, Sally Horne-Badovinac, Kristin Sherrard, Dan Kiehart, Lauren Anlo, Maria Bustillo, and Jessica Treisman for sending fly lines and antibodies and helpful conversations, Nat Prunet and the Biology Imaging Core for technical support, Steve Rogers, Vicky Bautch, and Dan Bergstralh for helpful feedback on the manuscript, and the Peifer, Bergstralh/Finegan and Williams labs for feedback throughout. M.C.B was supported by the DFG Walter Benjamin Programme (ref. GZ: BI 2384/1-1).and work in the Peifer lab is supported by NIH R35 GM118096.

## Supplement

**Supplemental Table1: Screen for Semaphorins in all adjacent cells**

**Supplemental Results 1: Semaphorin1B acts in TNM to maintain muscle coverage and is the only Semaphorin whose reduction affects this process**

We next asked which Semphorin ligands are relevant in TNM and whether they act from adjacent tissues or at TNM-TNM contacts. PlexA binds the transmembrane Semaphorins Sema1a, Sema1b, and Sema5c while Sema2a and Sema2b bind PlexB[74]. To identify the binding partners of PlexA that are relevant in vivo in TNM, we knocked down all known Semaphorins in each adjacent tissue. For expression in TNM, we used our driver specific to muscles (*dMef2-Gal4*). RNAi of *sema1b* led to defects in muscle coverage while RNAi of *sema1a* and *sema5c* did not (Supplemental Table 1, Fig 1e). One of the four *sema1a* lines caused defects in muscle shape at the base of the testis but these were completely distinct from the phenotypes we observed in *plexA*^RNAi^ or *sema1b*^RNAi^ (Fig S1c). Further, *sema1a^k13702^* – a mutant caused by P-element insertion into the 5’ UTR thought to result in complete loss of function[25] – does not cause any shape or coverage defects in adult testes (Fig S1d).

As there is no driver known to us that is exclusively expressed in pigment cells, we used *htl-Gal4* that drives in <u>both</u> pigment cells and myotubes[11] to deduce if pigment cells influence myotube migration via Semaphorins. This driver yielded the same results as seen with *dMef2-Gal4* with no changes in phenotype intensity (Supplemental Table 1), which suggests there is no additional role for any of the Semaphorins in the pigment cells. We also used *six4-Gal4*, a driver that drives expression in all somatic testis cells, including the cyst cells that underlie the migrating TNM[75]. Using this driver none of the *semaphorin*^RNAi^ constructs caused defects (Supplemental Table 1). In summary, our data reveal that Sema1b has an important function in testis morphogenesis that is TNM-autonomous.

**Supplemental Results 2: Expression of *plexA*^RNAi^, *sema1b*^RNAi^ and PlexA^SA^ affects migration speed and distance.**

After *plexA*^RNAi^ mean short-term cell motility was slowed: 0.77±0.04 µm/5 mins after *plexA*^RNAi^ versus 0.94±0.07 µm/5 mins in wildtype. Total displacement on the proximodistal axis was also altered (Figure 4a, arrow). Wildtype cells move 55±19 µm over 400 mins, while *plexA*^RNAi^ cells only move 35±17 µm (Fig. 4b).

*sema1b*^RNAi^ also reduced mean short-term cell motility: *sema1b*^RNAi^ cells moved 0.74±0.06 µm/5 mins versus 0.94±0.12 µm/5 mins for controls. Long-term motility was also affected – wildtype cells moved 57±8.7 µm while *sema1b*^RNAi^ cells moved 41±5.7 µm (Fig. 4c). PlexA^SA^-overexpression too, reduced both short-term (PlexA^SA^ cells 0.74±0.09µm/5 mins; controls 0.92±0.09µm/5 mins) and long term motility (PlexA^SA^ cells 37±8 µm; controls 50±9 µm) (Fig. 4d).

**Supplemental Results 3: The role of PlexA in TNM is independent of Myosin-dependent gap-closure**

PlexA’s best-known function is in contact-dependent repulsion or defasiculation[24], which is typically connected to RhoA-dependent activation of Myosin-contractility[76]. Thus, we chose to examine Myosin dynamics upon knockdown. *myosin* knockdown causes muscle gap formation in *Drosophila* testes[11], roughly reminiscent of *plexA* and *sema1b* knockdown phenotypes. Previous data suggested that one role of Myosin during TNM migration lies in promoting contractility of supracellular actin cables that support gap closure during migration[10]. 4D over-night imaging (Video S7, Fig S6a, b) revealed that knockdown of non-muscle (NM) myosin2 light chain – called Spaghetti Squash (Sqh) in *Drosophila* – reduced supracellular cables surrounding gaps, and they were replaced by filopodia, reminiscent of wound closure[6].

We next examined whether *plexA*^RNAi^ altered Myosin localization or contractility. We first imaged wildtype TNM expressing NM-Myosin2-GFP under the control of *dMef2*-Gal4[77]. This revealed that non-muscle Myosin is organized into filaments that are evenly distributed in the cell (Fig S6c, Video S8). Overexpression also causes formation of large artificial clusters that don’t take part in the contractions and that are not present in cells expressing endogenously tagged versions of Sqh (*data not shown*). This myosin network causes cells to pulsate, and waves of contractility move across the cell body (Video S8). These behaviors are also apparent when imaging cells expressing the F-actin probe Lifeact::GFP (Video S9). The pulsation is absent from *NM-Myosin*^RNAi^ TNM (Video S9). In *plexA* knockdown TNM, myosin appears more condensed (Fig S6d), consistent with the general cell-shape changes. However, pulsation still occurs in areas in which cells are flat (Video S8), thus contrasting from the effects of myosin knockdown (*sqh*^RNAi^, Video S9). These data suggest that *plexA* knockdown does not simply reduce or eliminate myosin contractility. Further, the effects of *plexA*^RNAi^ and *NM-myosin*^RNAi^ (*sqh*^RNAi^) on muscle coverage are synergistic upon double knockdown, consistent with the idea that myosin’s roles in closing gaps in muscle coverage are independent from those of PlexA signaling (Fig S6e–g). *sema1b* ^RNAi^ also had synergistic effects when combined with *NM-myosin*^RNAi^. This suggests that Myosin and Plexin/Sempaphorin signaling are involved in two independent mechanisms that seal gaps during migration.

---

**Figure S1.**
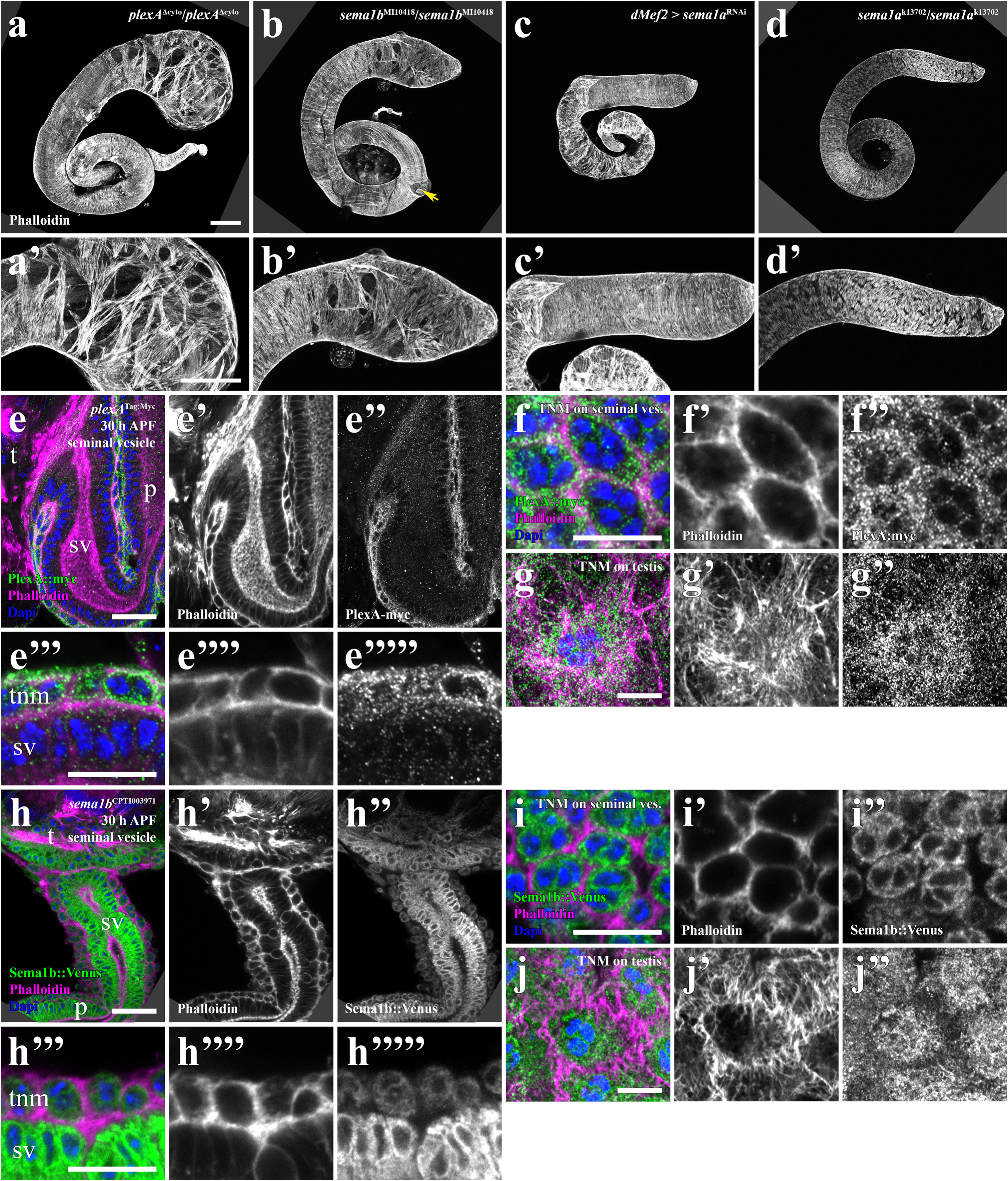
(a–d) Adult testis and magnification of the hub region in (a’–d’) in *plexA*^Δcyto^/*plexA*^Δcyto^ (a) and *sema1b*^MI10418^/ *sema1b*^MI10418^ (b), *sema1a*^RNAi^ (c) and *sema1a*^k13702^/ *sema1a*^k13702^ (d)-background. Musculature stained with phalloidin. Scale bars: 100 µm. (e–g’’) Airyscan 2 micrographs of fixed/mounted 30 h APF testis of a stock expressing an endogenously myc-tagged version of *plexA*. PlexA is strongly enriched in TNM with respect to other tissues in the testis. Phalloidin (magenta), Dapi (blue) and an anti-myc antibody (green). (e–e’’’’’, f–f’’) TNM on the seminal vesicle before migration. (g-g’’) TNM migrating on the testis. (h-j’’) Airyscan 2 micrographs of fixed/mounted testis of a line with Venus-tagged Sema1B (30 h APF at 26.5 °C) stained with Phalloidin (magenta), Dapi (blue) and an anti-GFP antibody (green). Stationary TNM on the seminal vesicle (h–h’’’’’, i–i’’) and migratory on the testis (j-j’’) are shown. (e–j’’) t=testis, sv=seminal vesicle, p=paragonium. (e–e‘‘, h–h‘‘) Scale bars= 20µm (e‘‘‘–g‘‘, h‘‘‘–j‘‘) Scale bars= 10µm

**Figure S2.**
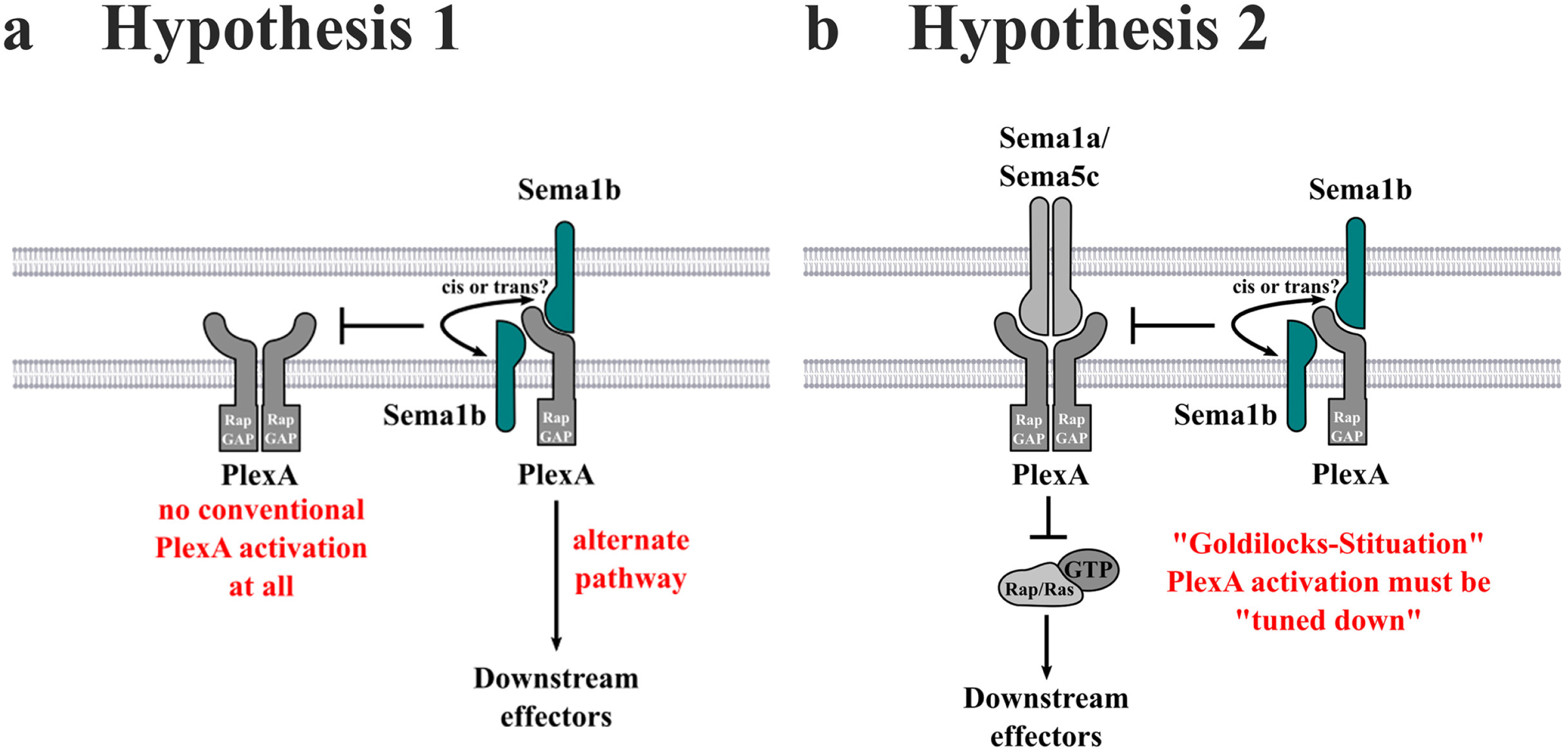
Schematic of potential ways PlexA and Sema1B might interact during TNM migration.

**Figure S3.**
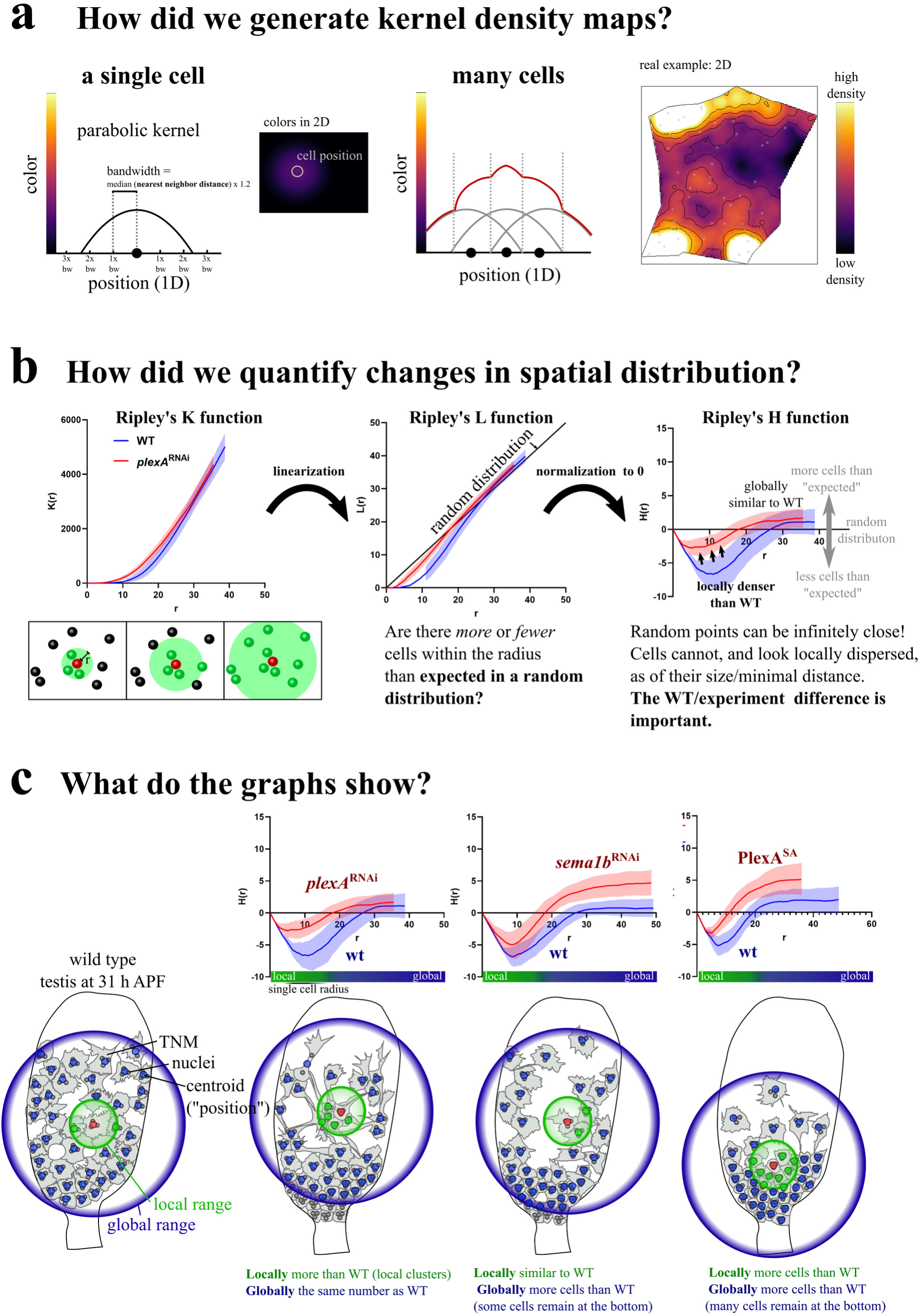
Related to Fig. 2 and 3. Detailed explanation of spatial statics used in this work. (a) For kernel distribution maps we used a parabola-shaped (*Epanechnikov*)-Kernel. To make local differences visible – even if density in general is very high – we coupled kernel bandwidth (sigma) to the median nearest-neighbor-distance. (left) Kernels are visible as color gradients around a cell position that is depicted on the y-axis. (right) If multiple cells are close to each other, values get added, resulting in a lighter color. (b) Ripley’s **K function** measures if points are clustered, random or dispersed. The measurement is based on the number of positions in a growing circle r. At each radius r, K(r) can be higher or lower than a theoretical random distribution (*point poisson process*). A higher value shows clustering, a lower one dispersal. The graph is easier to read after linearization to the **L-function**, in which L(r) = r (diagonal line) matches a random distribution. L-Function can be further transformed, so, that the “random-point-line” becomes the x-axis, resulting in the **H-function**. The example shows that both, WT and experiment are clustered when compared to “random points”. That’s because cell size does not allow cells to be infinitely close, which is possible for points in a *poisson process*. Because of TNM being relatively large when compared to the size of the total system, this effect has a strong influence. Nevertheless, in our biological application, it is the difference between WT and experiment that is important. This difference in this example shows that cells in the experiment are more clustered at small radii. (c) In wildtype cells are highly dispersed at low radii, which fits with what is visible in microscopic data (Fig 2a). On a larger scale, there is a certain level of clustering measured, consistent with the fact, that there are gaps in the cell sheet. *plexA*^RNAi^ cells are much more clustered (higher) at low radii. This is at a local level, close the size of a single WT cell. Globally, the number of cells compared to a random distribution is identical to WT, showing clustering to be exclusively a local effect. *sema1b*^RNAi^ shows the opposite trend. Globally, homogeneity is strongly affected, while the curves become closer to WT, the smaller the radii. This suggests a global effect on homogeneity, due to more distinct clusters and potentially a high density of cells at the base. This effect is similar but much stronger pronounced in PlexA^SA^ where we measured strong global clustering but also strong local densification, which fits to the observation that rear cells remain dense while front cells often completely lose contact.

**Figure S4.**
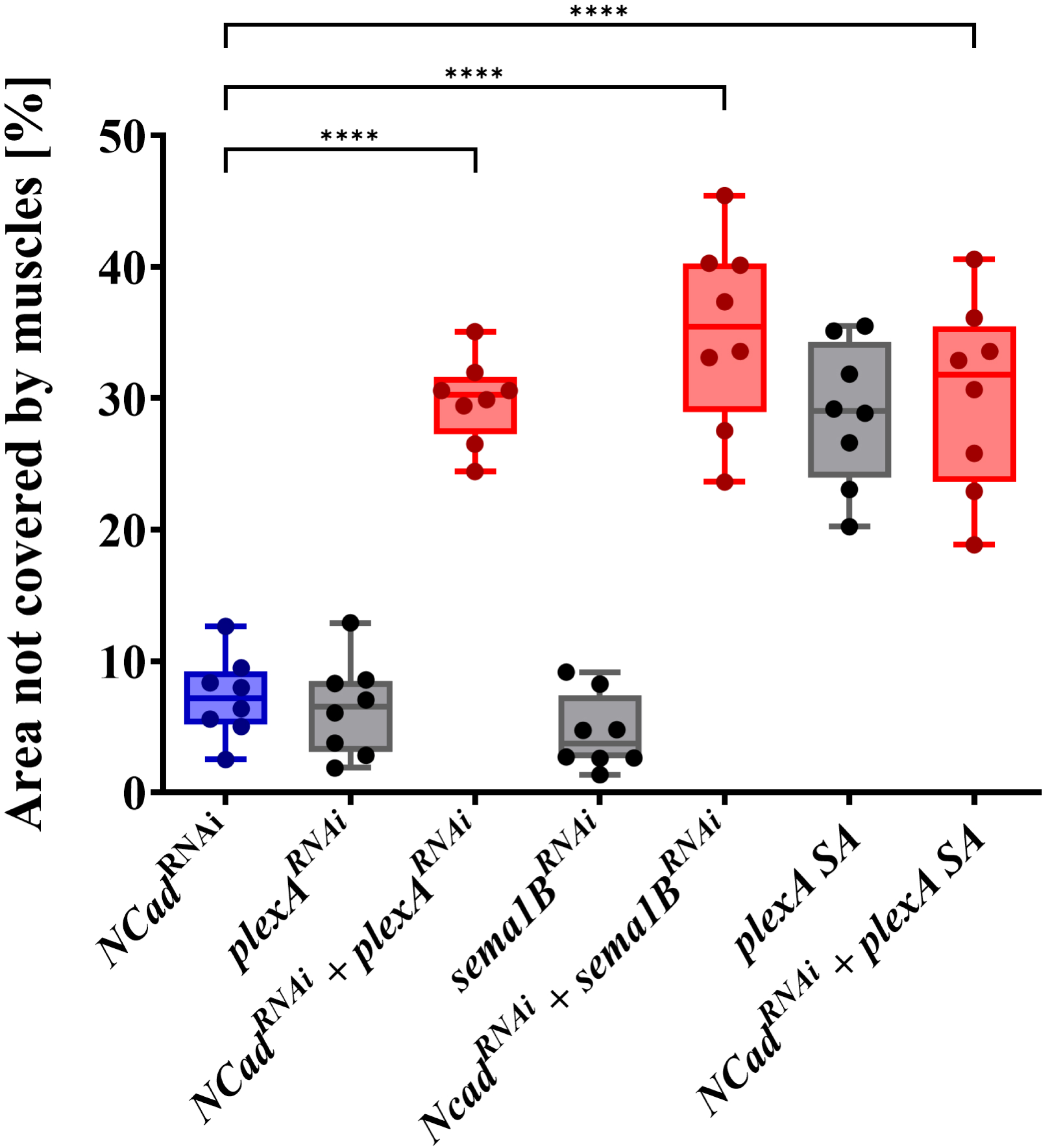
Related to Fig. 5. Quantification of area uncovered upon genetic interaction experiments with N-Cadherin. n=8 testes/condition. Statistical test: Ordinary one-way ANOVA, Šídák’s multiple comparison test. P-Values (all): < 0.0001.

**Figure S5.**
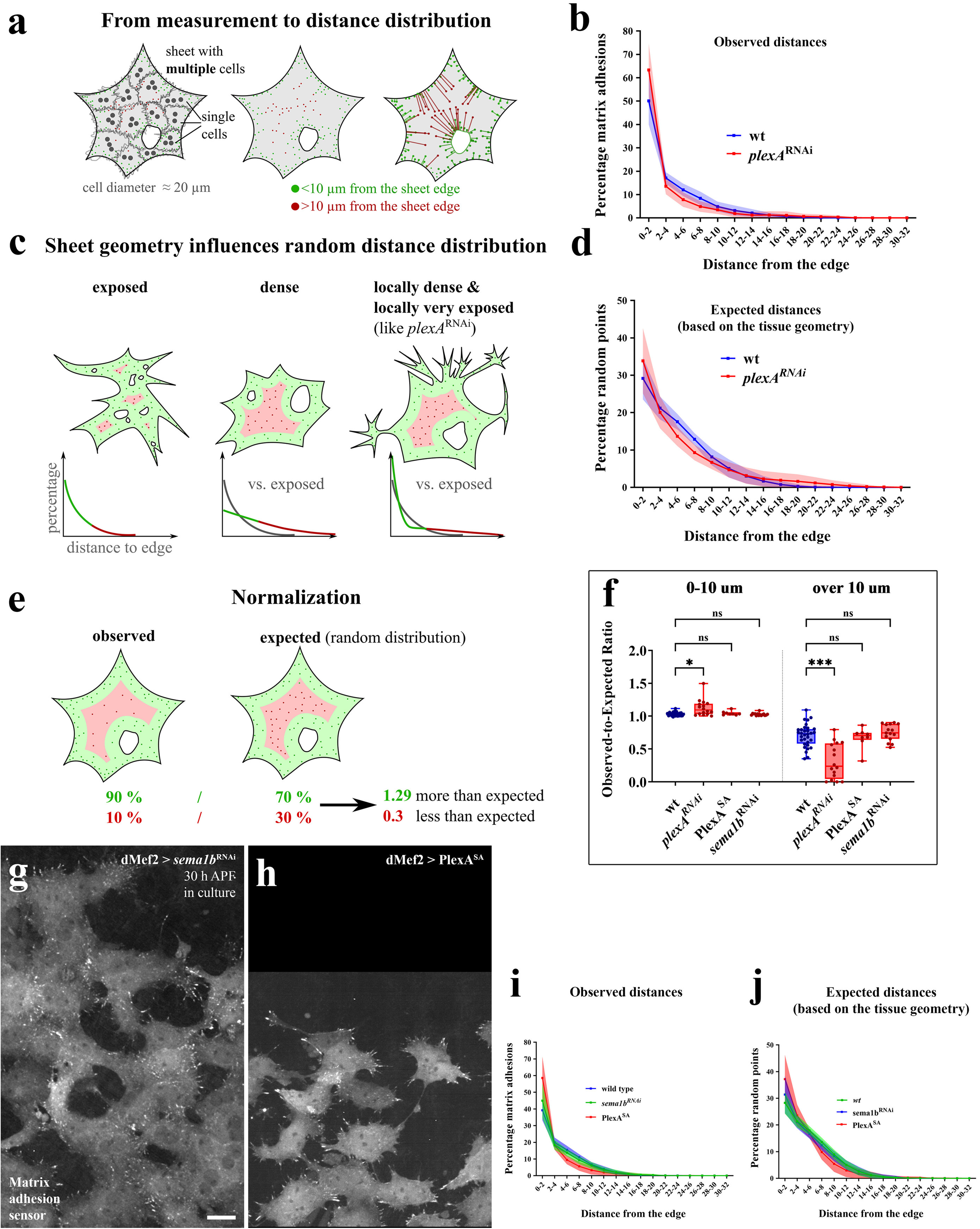
Detailed Explanation of the Quantification of Integrin adhesion distribution. Related to Fig. 6. (a) We quantified the position of integrin-based adhesion within cell sheets. Because we only know the outlines of the cell sheet – not the position of cell-cell borders – we quantified the distance between each matrix adhesion and the closest part of the free edge. Free edge includes also holes within the sheet. Adhesions that are closer that 10 µm are depicted in green, adhesions that are farther that 10 µm are depicted in red. In reality, we carried out the quantification in ROIs, with the edges of the ROI not being interpreted as sheet edges by the R-script. (b) This quantification (pictures in Fig. 6 a’’, b’’) revealed that plexA has much more adhesions very close to the free edge (0-2 µm), but much less farther adhesions. Although, in extreme remote parts of the cell, there are no more adhesions in wildtype but very few in *plexA*^RNAi^ because of differences in the tissue geometry elucidated below. n = 8 testes. Line = Mean of each individual distribution/testis. Shaded area = SD. (c) Even purely random points would have different length-distributions in different geometries. Very exposed geometries have few remote areas and therefore fewer remote points, than denser geometries. Geometries that have local dense areas, but also extremely exposed areas (like in *plexA*^RNAi^) have both, more close point – but also more remote points. (d) The quantification of random points in wildtype and plexA (pictures in Fig. 6 a’’’, b’’’) looks exactly as the hypothetical distribution in (c). plexA has more close but also more extremely remote random points compared to wildtype. This finding explains why in the curve in (b), there are extremely remote adhesions in *plexA*^RNAi^, absent from wt. This method was used for the subsequent normalization but is also a nice tool to quantify the tissue geometry in general. It is consistent with our observations that *plexA*^RNAi^ causes local densifications in the cell sheet, but also elongated processes and growth-cone like protrusions, causing locally exposed areas. (e) We used these data to normalize the data from (b). To simplify the quantification, we focussed on adhesions close to the cell edge (< 10 µm ≈ cell diameter/2) and remote adhesions (> 10 µm distance) and calculated their proportion. We calculated the same proportion for random points. Then, we calculated the observed/expected ratio for both, the close and the remote adhesion percentage. If the proportion for either is lower in the random distribution (<1), there are less than expected adhesions and if it’s higher (>1), there are more than expected. (f) Quantification from Fig. 6c. This quantification revealed that there are slightly more “close” adhesions but much less “remote” adhesions in plexA^RNAi^, consistent with an absence of cell-cell adjacent matrix adhesions while free edge matrix adhesions are normal. (g-j) High resolution life cell imaging of ex-vivo cultured (30 h APF at 26.5 °C) testis. TNM express *Focal adhesion targeting domain (fat)::eGFP* under the control of *dMef2-Gal4* in *sema1b*^RNAi^ (g) and *PlexA*^SA^ overexpression (h). Scale bar: 10 µm. Observed distances (i) and expected distances (j) where quantified as explained above. Consistent with the described cell-dispersion, both conditions show a distribution indicative of a more exposed but not locally densified geometry. n = 8 testes. Line = Mean of each individual distribution/testis. Shaded area = SD.

**Fig. S6.**
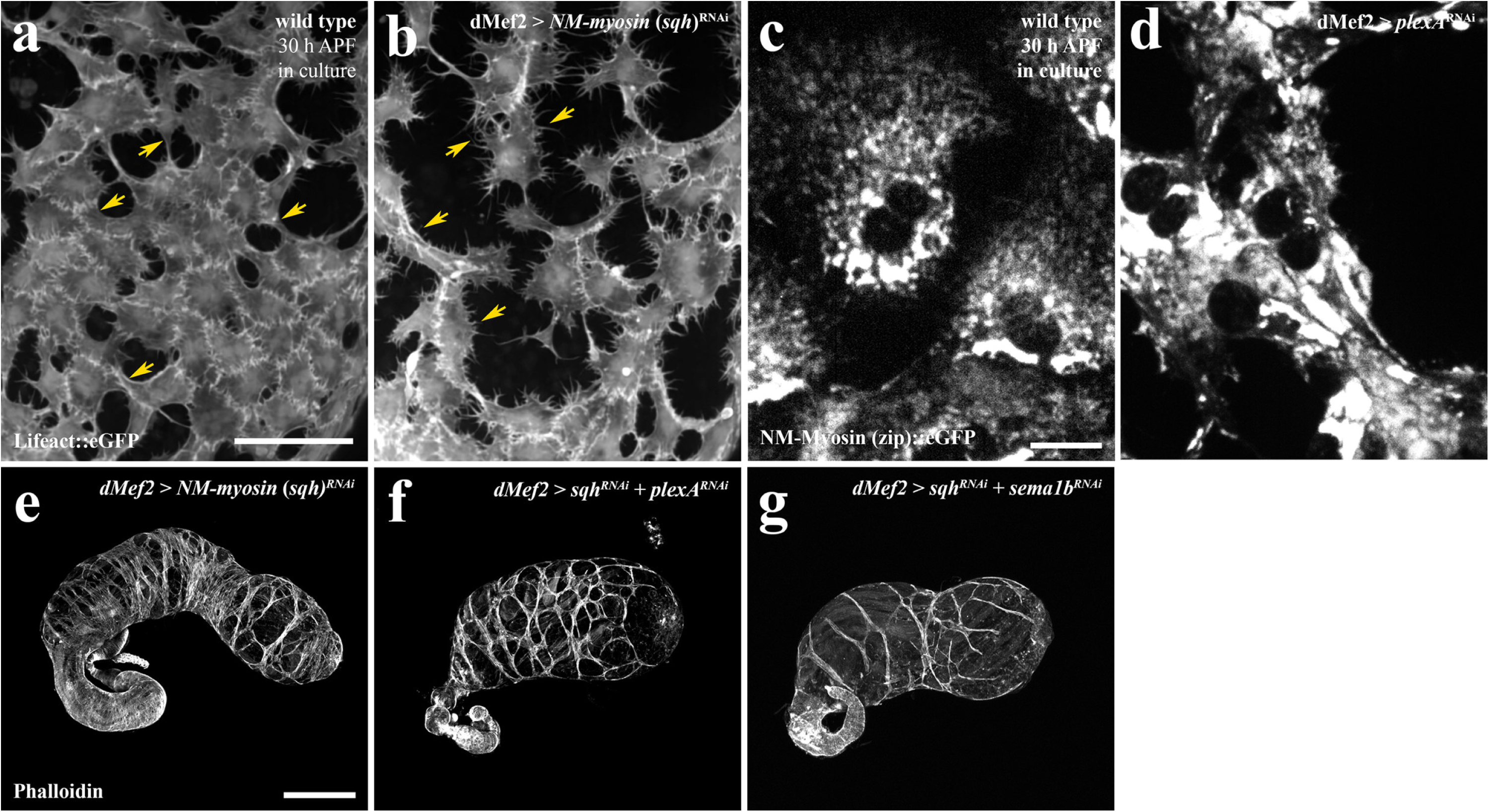
Myosin depletion causes gap-closure defects, but this process in independent of PlexA and Sema1B. (a,b) Ex-vivo culture of 30 h (26.5 °C) testis. TNM express *Lifeact::eGFP* and *Cherry^nls^* (not shown) under the control of *dMef2-Gal4*. Wildtype (a), *NM-Myosin*^RNAi^ (b). Yellow arrows (a): edges with cables. (b) edges with filopodia instead. Scale bar = 100 µm. (c,d) Ex-vivo culture of 30 h (26.5 °C) testis with TNM expressing UAS-NM-myosin (zip)::eGFP under the control of *dMef2-Gal4*. (e–g) Genetic interaction study of *NM-myosin (sqh)* (e) with *plexA* (f) and *sema1b* (g). Adult testes were stained with Phalloidin, Scale bar: 100 µm.

## Video Legends

**Video S1.** Overview of TNM migration in culture of a wild type testis 30 h APF with TNM tracked using Imaris. Timesteps: 5 min. Lifact::eGFP marks actin and Cherry^nls^ nuclei. (used for tracking nuclei).

**Video S2.** TNM migration in cultured testes in wild type background, plexA^RNAi^ background and sema1b^RNAi^ background. Lifact::eGFP marks actin and Cherry^nls^ nuclei. 30 h APF. Timesteps: 5 min.

**Video S3**. High resolutions videos of TNM type background and in plexA^RNAi^ background. Lifact::eGFP marks actin. Timesteps: 20 sec.

**Video S4**. Example for “Closest Neighbor Quantification” using Imaris. Color code of the “balls” shows distance to the closest neigbor. plexA^RNAi^ background. 30 h APF. Timesteps: 5 min. Lifact::eGFP marks actin.

**Video S5**. Videos of Kernel Density Quantification. Details in Supp. Fig. S3.

**Video S6.** TNM migration in cultured testes in wild type background and in plexA^SA^ background. Lifact::eGFP marks actin and Cherry^nls^ nuclei. 30 h APF. Timesteps: 5 min.

**Video S7.** TNM migration in cultured testes in wild type background and in *sqh* (NM-myosin-RLC)^RNAi^ background. Lifact::eGFP marks actin and Cherry^nls^ nuclei. 30 h APF. Timesteps: 5 min.

**Video S8**. High resolutions videos of TNM type background and in plexA^RNAi^ background. zip::eGFP marks NM-myosin. Timesteps: 20 sec.

**Video S9**. High resolutions videos of TNM type background and in sqh^RNAi^ background. Lifact::eGFP marks actin. Timesteps: 20 sec.

## Notes

### Competing Interest Statement

The authors have declared no competing interest.

