## Supplementary figures and images for "Plexin/Semaphorin Antagonism Orchestrates Collective Cell Migration, Gap Closure and Organ sculpting by Contact-Mesenchymalization"

### Supplemental Figure 1

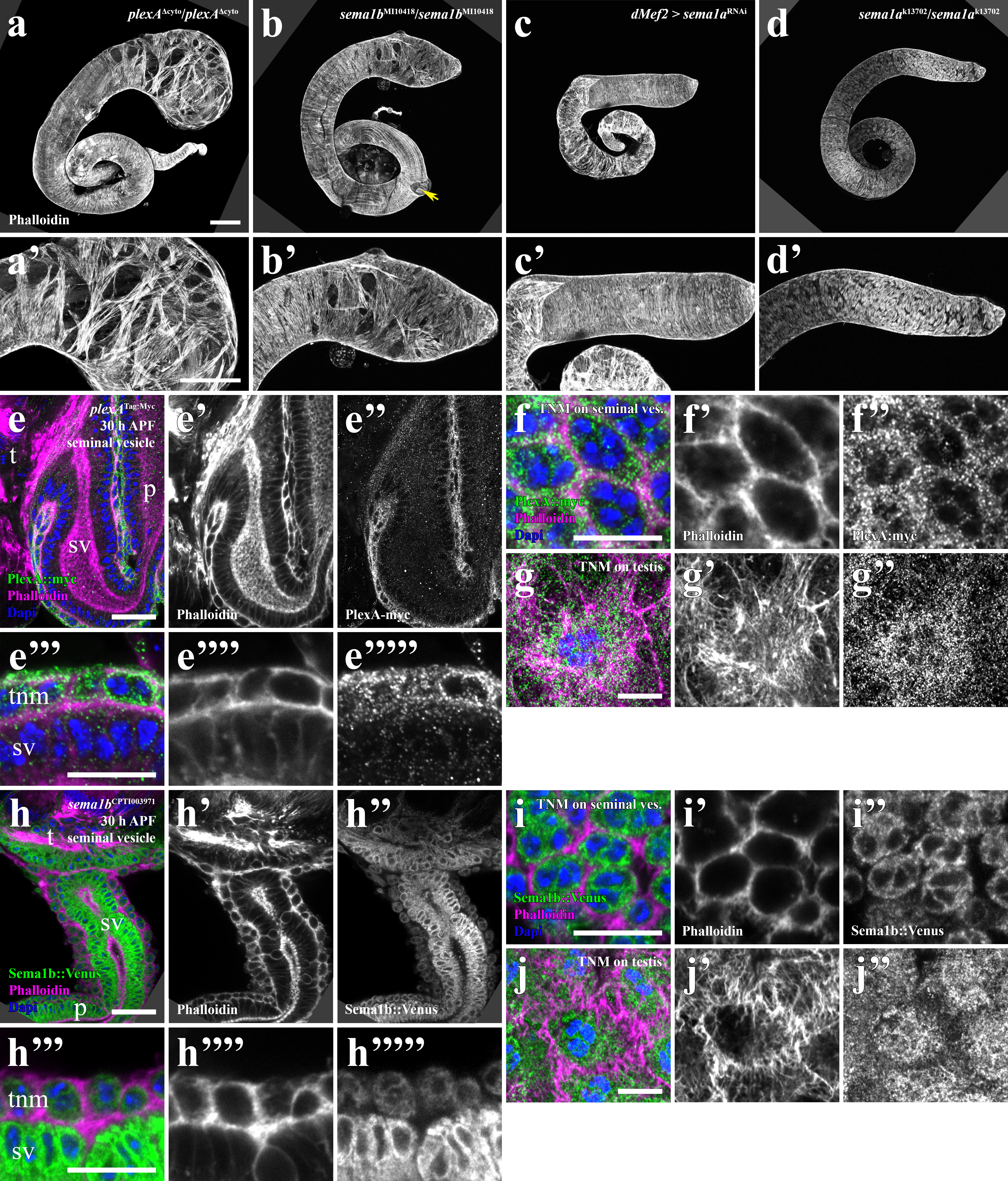

### Supplemental Figure 2

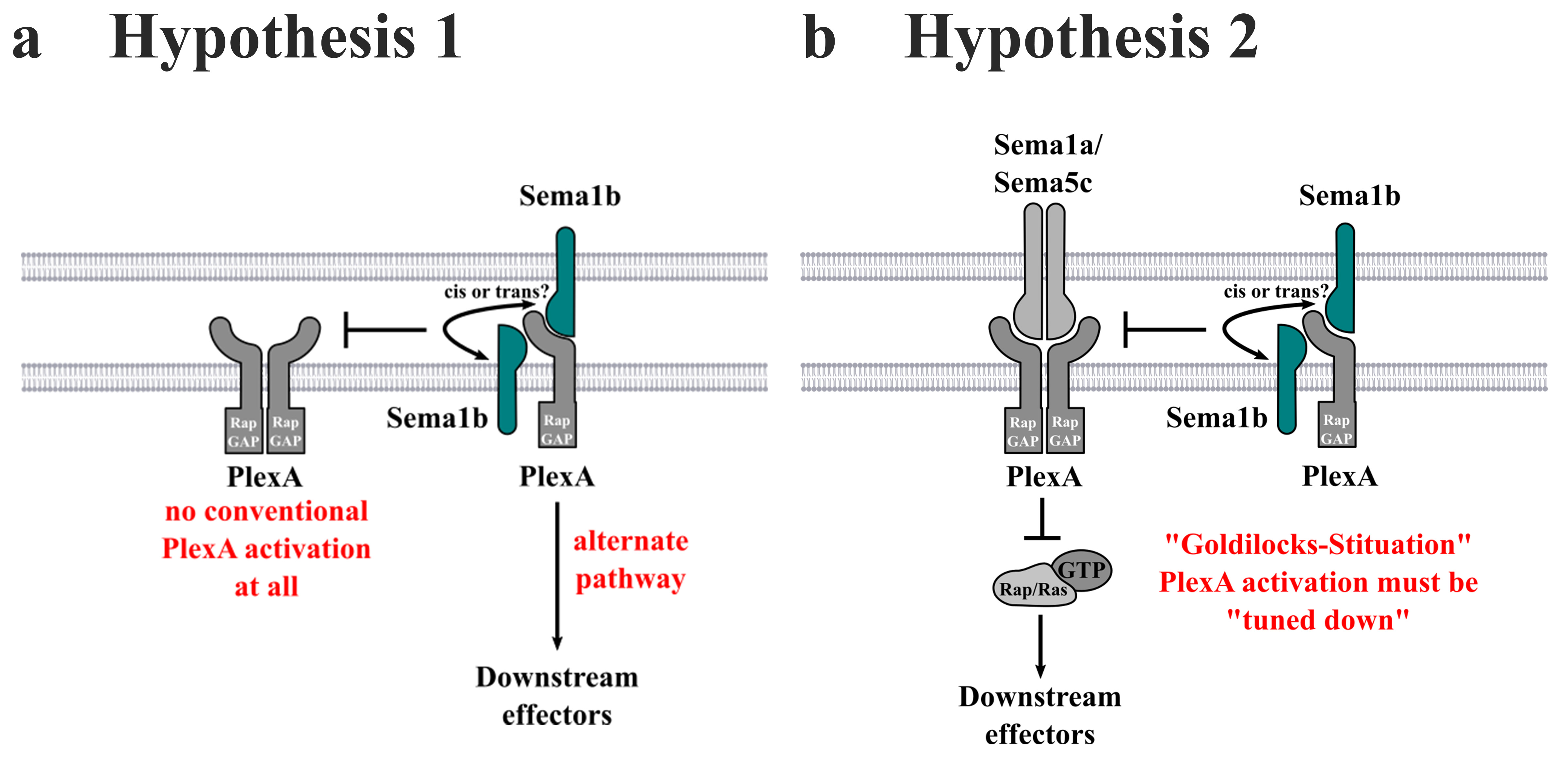

### Supplemental Figure 3

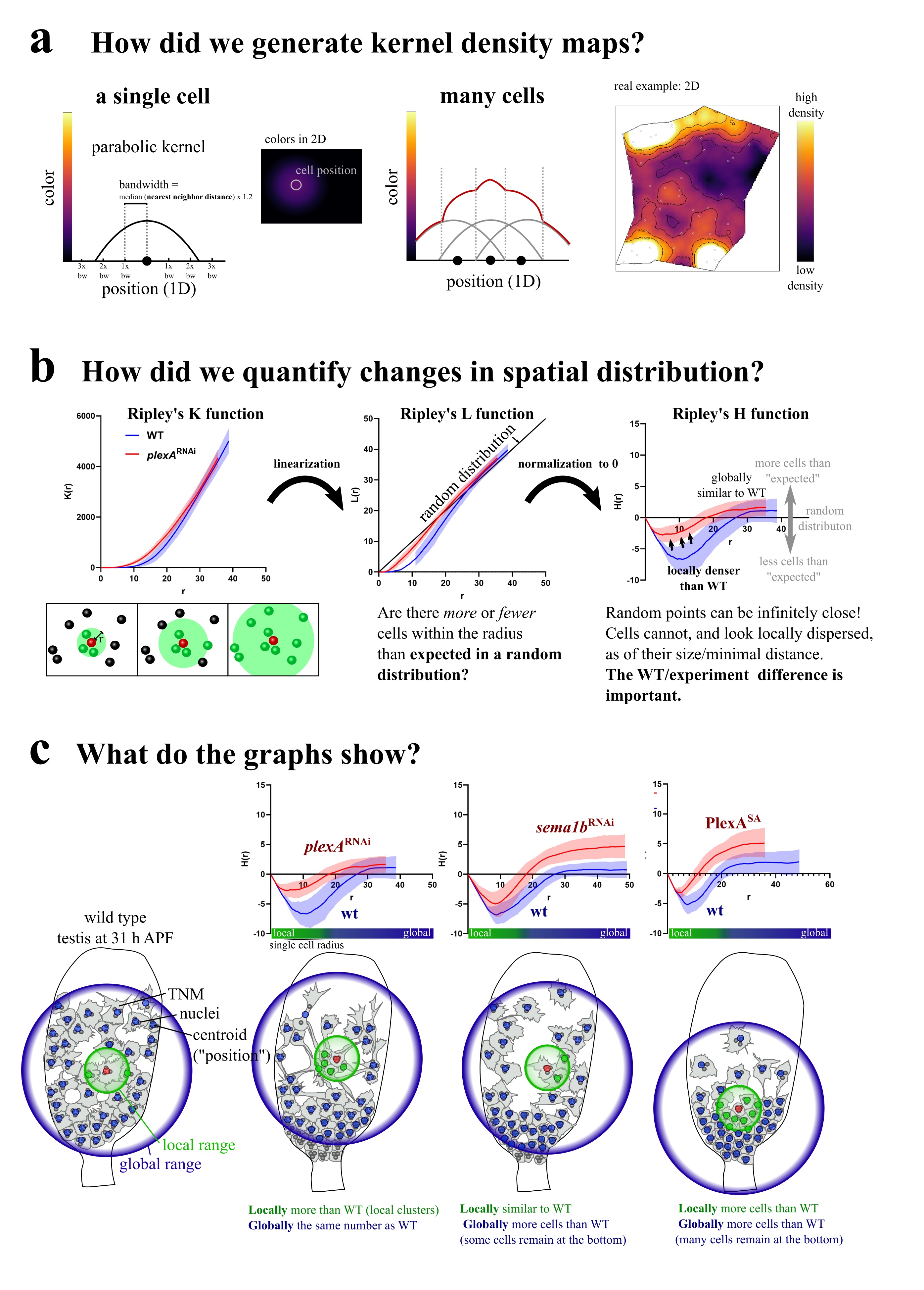

### Supplemental Figure 4

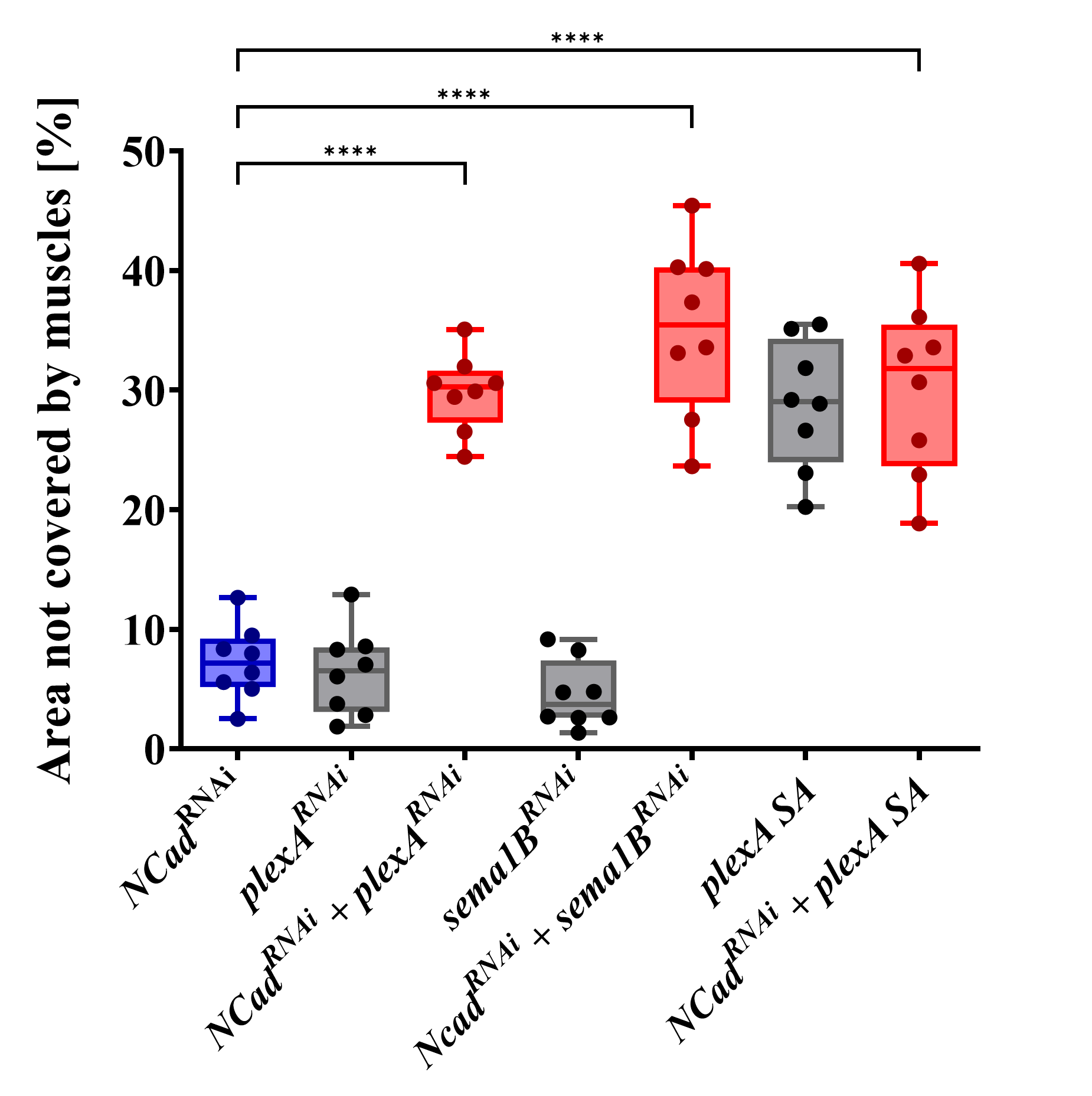

### Supplemental Figure 5

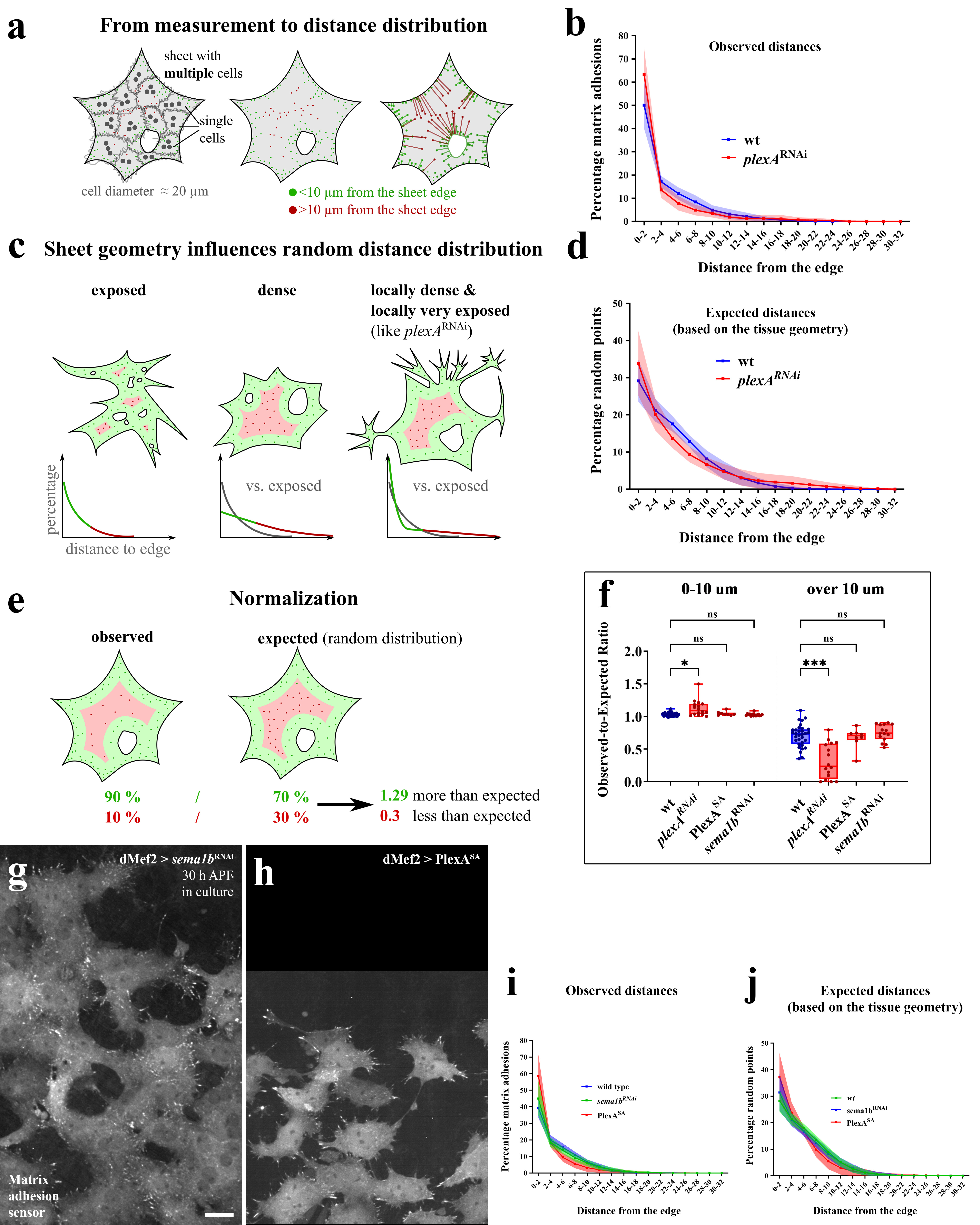

### Supplemental Figure 6

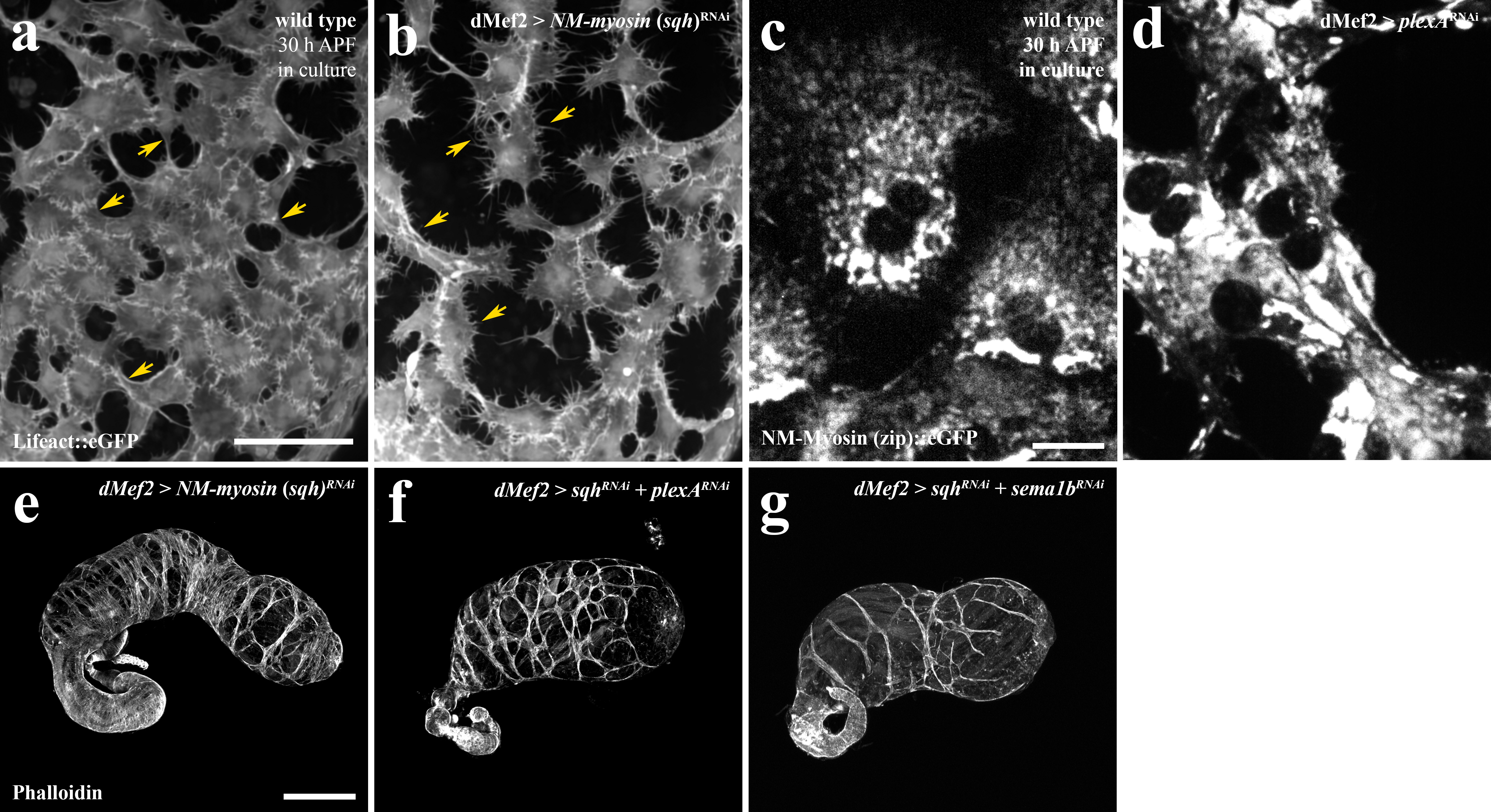
